# Clustered Inputs Maximize Efficiency for Stable Place Field Encoding in Hippocampal Pyramidal Neurons

**DOI:** 10.1101/2025.05.30.657022

**Authors:** Simone Tasciotti, Daniel Maxim Iascone, Spyridon Chavlis, Luke Hammond, Yardena Katz, Attila Losonczy, Franck Polleux, Panayiota Poirazi

**Author notes:** Present address: Department of Neurology, The Ohio State University, Wexner Medical School, Columbus, Ohio, USA. Present address: O’Donnell Brain Institute at UT Southwestern Medical Center, Harry Hines Blvd. Dallas, Texas, USA.

## Abstract

How spatial organization of synaptic inputs along the dendritic tree of neurons influence their feature selectivity is a central question in neuroscience. Here, we mapped the three-dimensional distribution of all excitatory and inhibitory synapses across the entire dendritic arbor of individual mouse CA1 pyramidal neurons in vivo and built biophysically detailed computational models to probe their functional impact on their ability to emerge as place cells. We found that excitatory, but not inhibitory, synapses are non-uniformly distributed, forming structural clusters preferentially on terminal dendrites. These excitatory synaptic clusters generate high-quality place fields more efficiently than randomized synaptic distributions, requiring fewer active synapses to achieve equivalent somatic output. Crucially, even when firing rates are matched, clustered inputs sustain significantly higher voltage-gated calcium influx and NMDA receptor activation, key substrates for synaptic plasticity. Further analysis reveals that clustering enables domain-specific computational strategies: oblique dendrites rely on cluster location, basal dendrites on cumulative synaptic strength, and the trunk on local input dispersion. Disrupting clustering collapses this compartmentalized processing into uniform summation. Our results establish synaptic clustering as a key mechanism that maximizes computational efficiency and enables sophisticated dendritic processing underlying hippocampal spatial representation.

## Introduction

The hippocampus and particularly its CA1 region plays a crucial role in spatial navigation. CA1 pyramidal neurons (PNs), specifically the spatially tuned ‘place cells’, are widely hypothesized to construct cognitive maps of the environment by combining spatial information and contextual cues^1–3^. Studies have shown that place cells are highly dynamic, undergoing remapping when an animal encounters a new or altered environment^2,4^. During remapping, CA1 place cells reorganize their spatial firing patterns in response to environmental changes, effectively encoding a new cognitive map^5^. In familiar settings, place cells exhibit stable firing fields to support the retrieval of established memories, while in novel contexts, rapid remapping enables the encoding of new spatial information.

This flexibility stems from the integrative properties of CA1 PNs, which allow them to adjust the salience of specific input streams based on the environmental context^6,7^. However, the electrophysiological and structural variability of these neurons hinders our in-depth understanding of their computational capabilities. Numerous studies have attempted to characterize how these properties, along with the organization of synaptic inputs, collectively shape the input-output dynamics of CA1 PNs^8–14^. Yet, the impact of the spatial distribution of excitatory (E) and inhibitory (I) synaptic inputs on the formation and functional features of place cells remains largely unexplored.

Synaptic organization along the dendritic arbor has been proposed to influence the integrative properties of specific classes of neurons. For example, projections from CA3 PNs predominantly target proximal dendrites, basal and apical obliques, while long-range inputs from the entorhinal cortex (EC) innervate the distal apical tuft dendrites of CA1 PNs^15,16^. This anatomical segregation allows both the independent processing of distinct information streams and their conditional integration, thus dynamically shaping the feature selectivity of CA1 PNs^17–19^. Additionally, differential ion channel distributions and morphological features of dendrites across these CA1 layers modulate synaptic efficacy, such that identical inputs can have variable impacts on neuronal output depending on their dendritic location^20,21^. A comprehensive exploration of how different incoming inputs shape the local dendritic properties is essential for elucidating the mechanisms underlying neuronal information processing.

Anatomical work revealed that rat CA1 PNs are characterized by non-uniform synaptic distributions at the dendritic branch level^9^. This type of synaptic organization can greatly affect the dendritic input-output function^22,23^. For example, synchronous activation of synapses within the same branch was shown to drive strong nonlinear dendritic responses^24–26^, a feature that generalizes to other cell types^27–29^. These nonlinear dendritic responses were eliminated when the same synapses were distributed across different dendrites^24,26^, highlighting the importance of synaptic distribution on CA1 input-output mapping.

The abovementioned effects are underlined by voltage-dependent ionic conductances in dendrites, which enable the induction of local regenerative events, turning neuron dendritic tree into a complex processing system with enhanced computational capabilities^22,24,26,30–32^. As the spatio-temporal organization of synaptic inputs is essential for triggering these local events, synapse location matters crucially^24,26,32^. Localized within dendritic compartments that host numerous nonlinear mechanisms, synapses engage in dynamic interactions, amplifying, attenuating, or gating signals based on their dendritic positions. This suggests that the formation and maintenance of synaptic contacts are not random processes but rather actively regulated mechanisms^33–38^. Recent findings further reveal that co-tuned inputs often converge onto spatially clustered synapses, creating what are known as dendritic “hotspots” for synaptic plasticity^39,40^. These hotspots can more easily produce non-linear dendritic events^41,42^, as suggested by earlier theoretical work^26,32,43^. Therefore, it is clear that to fully understand how CA1 PNs form place fields, it is essential to consider the spatiotemporal organization of their incoming inputs.

In this study, we mapped the precise spatial distribution of excitatory and inhibitory synapses in the entire dendritic tree of single mouse CA1 pyramidal neurons in order to determine how their organization shapes spatial tuning. Our reconstructions revealed that excitatory synapses are non-uniformly distributed, clustering significantly within terminal dendrites. Using validated biophysical models, we demonstrate that this clustered organization is far more efficient in forming high quality place fields than a randomized distribution, requiring approximately 13% fewer activated synapses.

Moreover, we found that the neuron’s computational strategy is dictated by its input architecture. Compared to randomized distribution, the biologically observed clustered configuration allows CA1 PNs to leverage distinct features associated with the different dendritic domains and drive somatic spiking. Conversely, when inputs are shuffled, CA1 PNs shift to a uniform strategy of cumulative, distributed summation across the dendritic tree. This suggests that synaptic clustering in CA1 pyramidal neurons is not merely a structural feature, but a functional mechanism for ensuring resource-efficient neural computation in the hippocampus.

## Results

### Reconstruction of CA1 Pyramidal Neuron Morphology

To reconstruct the dendritic morphology and synaptic distribution of all excitatory and inhibitory synapses of mouse dorsal CA1 PNs at single-cell resolution *in vivo*, we deployed a combination of sparse reporter expression, imaging, and computational tracing methods that we had previously developed^44^. We labeled dorsal CA1 PNs through *in utero* electroporation of plasmids expressing Cre-dependent Flex-tdTomato (red cell filler) and Flex-EGFP-Gephyrin (inhibitory synaptic marker) along with low levels of a Cre recombinase plasmid to achieve sparse labeling^45,46^ (**Figure 1A**). Using high-resolution confocal imaging (**Figure 1B**), combined with the open-source reconstruction software Vaa3D^47^, we first traced the entire dendritic tree (**Figure 1C**). Through the Vaa3D Synapse Detector toolkit^44^, we annotated all excitatory and inhibitory synapses along the entire dendritic tree. Given that the vast majority of excitatory synapses in CA1 PNs occur on dendritic spines^9^, we used them as a structural proxy of all excitatory synapses received by these neurons. Inhibitory synapses were identified by co-localizing EGFP-Gephyrin puncta with cytosolic tdTomato (**Figure 1B, Video S1**) as previously validated^44,48^. Using this method, we obtained detailed 3D mapping of each synapse’s position and shape, including features like synaptic volume, spine head diameter, and spine neck length. We also identified the relative location of inhibitory synapses, whether they are located on the dendritic shaft or on a dually innervated spine^49,50^. Finally, Vaa3D algorithms linked all synaptic features to the corresponding branch segment node, and neuron trace fragments of each CA1 PN from serial tissue sections were stitched together to create complete morphological reconstructions. This method resulted in full morphological reconstructions of the dendritic arbor along with a detailed 3D information about the position and size of each excitatory (**Figure 1D, Video S2**) and inhibitory (**Figure 1E, Video S3**) synapses for six CA1 PNs, including one that lacks the inhibitory synapse map. After implementing the synaptic reconstruction pipeline, we applied a custom smoothing algorithm to the dendritic diameter measurements to eliminate minor artifacts introduced by the reconstruction software (**see Methods, Figure S1**). From the structural analysis of these neurons, we observed, on average, 11,796 ± 2,212 (mean ± std, n=6 neurons, std: standard deviation) spines and 741 ± 545 inhibitory synapses, distributed over 7,164 ± 982 µm of average total dendritic length. Consistent with previous studies^51^, the average synaptic density, calculated as synapse/µm, was estimated to be 1.41 ± 0.18 for spines and 0.10 ± 0.07 for inhibitory synapses^52^ (**Figure 1F**).

**Figure 1:**
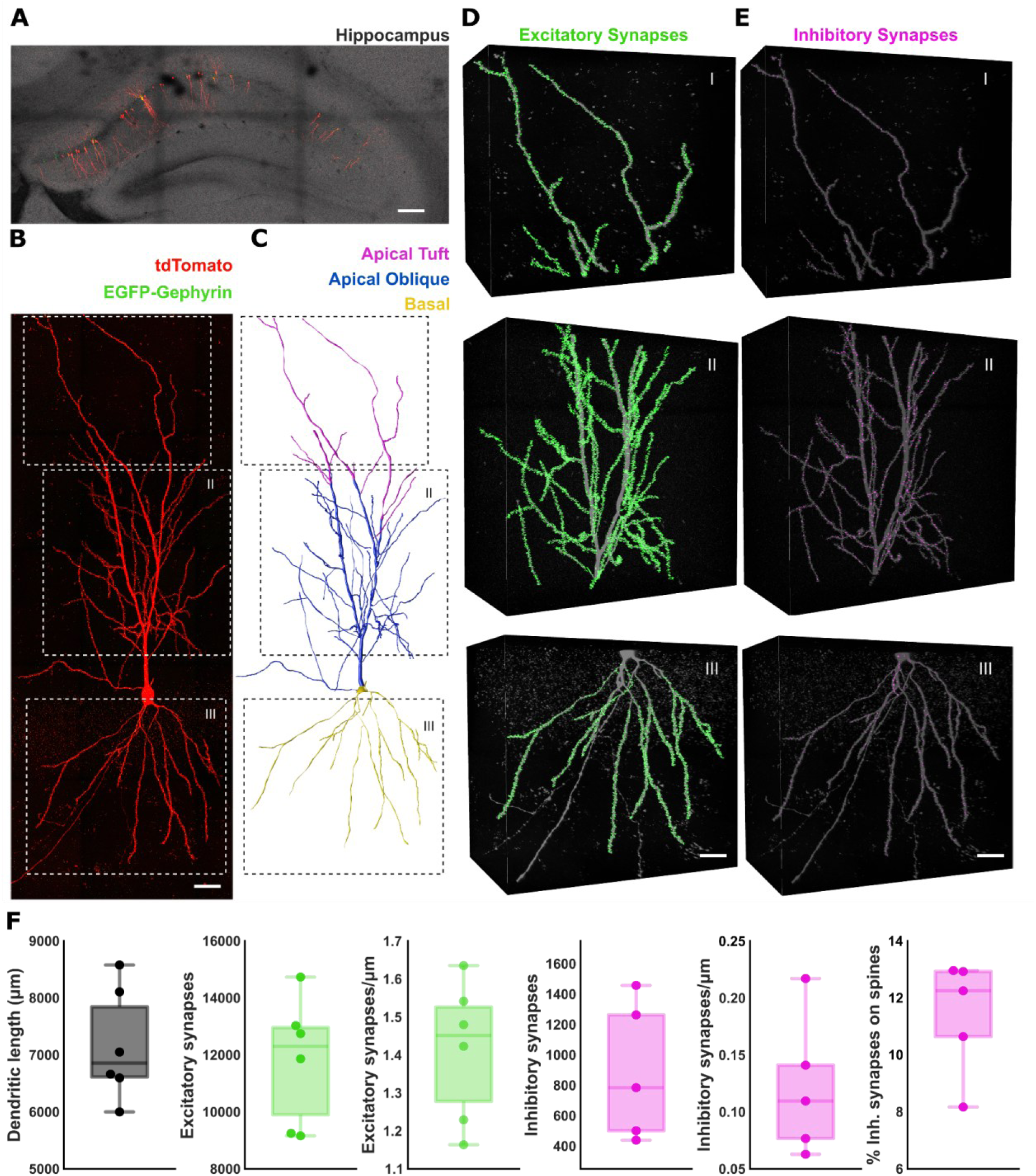
Reconstruction and analysis of all excitatory and inhibitory synapses across CA1 pyramidal neurons. **A**. Sparsely labeled pyramidal neurons expressing tdTomato (red) and EGFP-Gephyrin (green) in hippocampal CA1. Grey: tissue autofluorescence using 488nm excitation. Scale bar: 300 μm. **B**. Single-synapse resolution image of a CA1 pyramidal neuron expressing tdTomato (red) and EGFP-Gephyrin (green). Scale bar: 60 μm. **C**. Dendritic arbor of whole CA1 PN shown in panel B, annotated to highlight apical tuft dendrites (magenta), apical oblique and apical trunk dendrites (blue) and basal dendrites (yellow). **D-E**. All dendritic spines (excitatory synapses; D, green) and Gephyrin+ inhibitory synapses (E, purple) annotated throughout the dendritic arbor (grey) of one example reconstructed CA1 PN within insets from (B-C). Scale bars: 50 μm. **F.** Quantitative analysis of all reconstructed CA1 PNs showing dendritic structure and synaptic distribution. From left to right: total dendritic length (µm); total number of excitatory synapses and their density (syn/µm); total number of inhibitory synapses and their density (syn/µm); percentage of inhibitory synapses localized on a spine (dually innervated spines).

### Domain-Specific Analysis of Synaptic Organization in CA1 PNs

CA1 PNs exhibit significant variability in the structural and electrophysiological properties of their dendritic arbors, which are also targeted by different presynaptic sources which are remarkably segregated: (1) inputs from axons of CA2/CA3 PNs onto the basal dendrites, (2) inputs from axons from CA3 PNs only to the apical obliques dendrites and (3) inputs from axons from the entorhinal cortex (EC) on the apical tuft of CA1 PNs^8,9,53,54^. Consequently, we opted to analyze the synaptic organization by subdividing the dendrites into four distinct domains, namely *Basal*, *Apical Trunk*, *Oblique*, and *Tuft* dendritic domains. Within each domain, we further distinguished intermediate and terminal branches based on their relative branch order, referred to as *subdomains* (**Figure 2A**). We found that peri-somatic and intermediate dendritic regions exhibit a substantially lower spine density than terminal branches (**Figure 2B**), which is in line with previous findings^51^ while inhibitory synapses show no significant variation across subdomains (**Figure 2B**).

**Figure 2:**
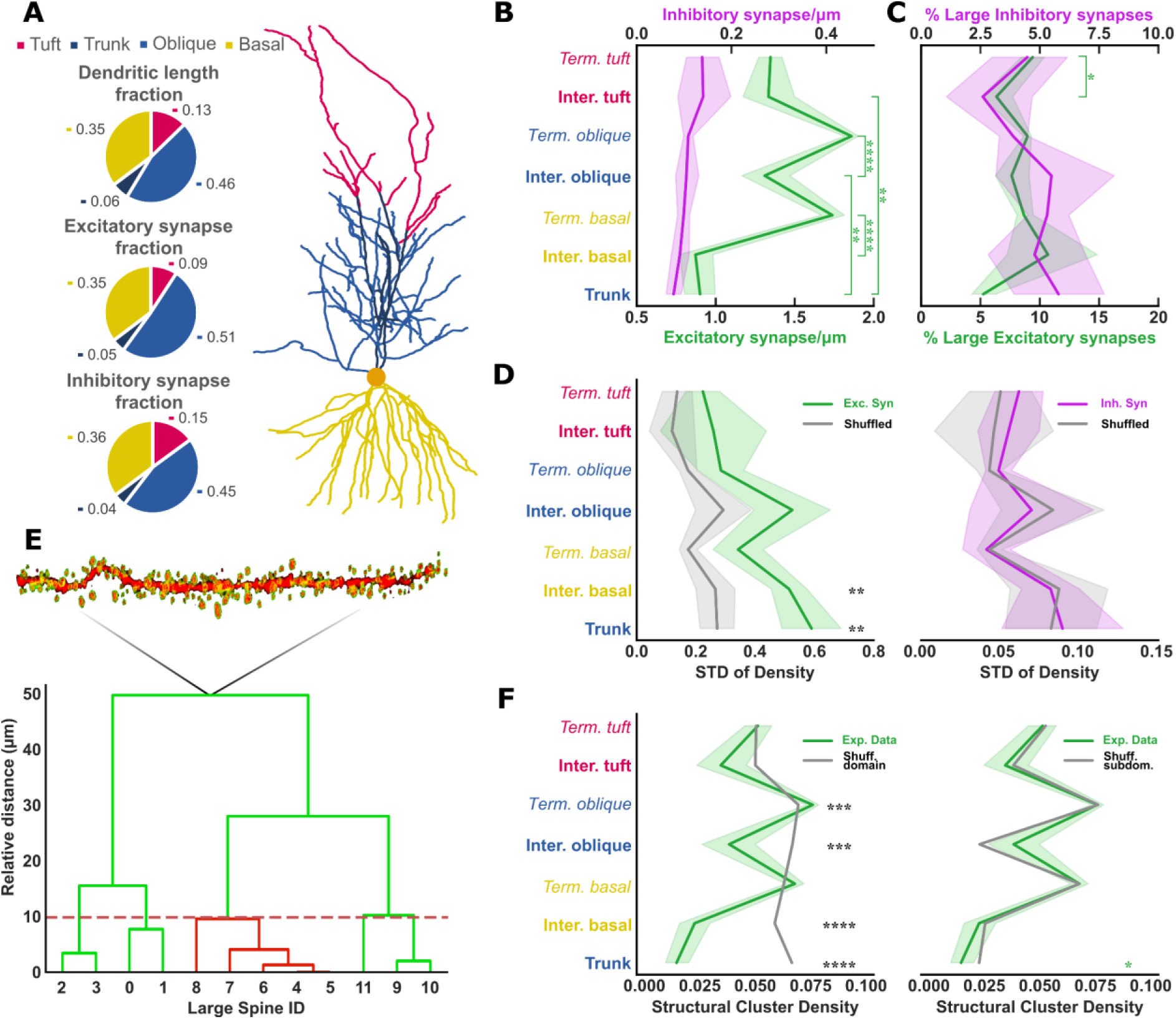
Analysis of Dendritic Subdomains and Synaptic Distributions. **A.** Left: pie charts showing the percentage of excitatory (E) and inhibitory (I) synapses located in different domains. Right: exemplary representation of a CA1 PN, with distinct dendritic domains highlighted in different colors. These include the Basal (yellow), Tuft (pink), Trunk (dark blue) and Oblique (light blue). Domains were further subdivided into intermediate and terminal subdomains, reported in the following panels with bold and italic, respectively. Terminal and Intermediate have been shortened to Term. and Inter. in figure for brevity. **B.** Density of E and I synapses across different subdomains. **C.** Percentage of large spines (i.e., spines with a volume above the 80^th^ percentile of the size distribution) for each subdomain. **D.** Variation (standard deviation, STD) in E and I synaptic densities for each subdomain. Experimentally observed STDs (green/purple) are compared to those expected from a random redistribution of synapses within their specific subdomains (grey). Specifically, synapses were randomly shuffled across branches within the same subdomain, and excitatory/inhibitory synaptic density variation was recalculated. Excitatory but not inhibitory synapse variability is significantly different from the shuffled distribution (2-way ANOVA for spine density std: distribution: F(1, 4228)=374.74, p<10^−10^; domain: F(6, 4228)=794.07, p<10^−10^; distribution x domain: F(6, 4228)=10.66, p<10^−10^). **E.** Schematic illustration of the hierarchical clustering algorithm used to identify spine clusters. The algorithm detects groups of large spines (with synaptic volumes above the 80th percentile) that are located within 10 micrometers of each other. The dashed line indicates the 10 μm distance cutoff. Red dendrogram strokes represent a spine cluster. **F.** Density of spine clusters across subdomains. Left: experimentally observed distribution (green) vs. a domain-specific shuffled distribution (grey). Right: experimentally observed distribution vs. a branch-wise shuffled distribution. Statistically significant differences are found only between the experimentally observed distribution and the shuffled-per-domain distribution. (2-way ANOVA for cluster density between ‘Exp. Dat’ and ‘Shuffled per domain’: protocol: F(1, 775761)=136.83, p<10^−10^; domain: F(6, 775761) = 3412.82, p<10^−10^; protocol x domain: F(6, 775761)=58.33, p<10^−10^). When shuffling within subdomains, the two distributions substantially overlap, except for the Trunk (2-way ANOVA for cluster density between ‘Exp. Dat’ and ‘Shuffled per subdomain’: protocol: F(1, 775761)=1.06, p=0.303; domain: F(6, 775761)=82383.08, p<10^-6^; protocol x domain: F(6, 775761) = 4.58, p = 1.19e-04). Pair-wise two-sided t-tests are shown in all plots. ^∗^p < 0.05, ^∗∗^p < 0.005, ^∗∗∗^p < 0.001, and ^∗∗∗∗^p < 0.0001, corrected for multiple comparisons using Bonferroni’s method. Error-bands in **B**, **C**, **D**, and **F** represent 95% confidence interval (CI) of the mean.

We also characterized the size of synapses in order to infer the degree of potentiation as a function of the dendritic subdomain. Synaptic volume corresponds to spine head size or postsynaptic density (PSD) size for excitatory and inhibitory synapses, respectively, and is linearly correlated with synaptic strength^22,55,56^. As previously described for layer 2/3 PNs in the mouse primary somatosensory cortex^44^, we used the Synapse Detector tool to map the distribution of synaptic strengths across all different subdomains. Large synapses were defined as those exceeding the 80^th^ percentile of the synaptic volume distribution of each cell, corresponding to the ∼70% increase in volume observed experimentally following LTP-inducing stimulation^57^. We found similar distributions of large synapses across dendritic subdomains, with the exception of the terminal Tuft, which exhibited a greater percentage of large spines compared to its intermediate counterpart (**Figure 2C**).

Another structural feature that could have important functional consequences is the branch-specific increase in synaptic density. During learning, new spines tend to form in proximity to existing spines, or appear in small clusters, demonstrating greater stability and marking clustered potentiation at the branch level^38,40,58–61^. To assess this possibility, we compared the experimentally observed variation (standard deviation, in short STD) in E and I synaptic density within dendritic subdomains to the synaptic densities of a subdomain-wise null distribution (**see Methods**) for each CA1 PN reconstructed. Dendritic subdomains with STD higher than that of the null distribution exhibit this branch-specific enrichment. We found branch-specific enrichment of excitatory synapses in the Trunk and intermediate Basal domains, whereas inhibitory synapses were overall uniformly distributed (**Figure 2D**). These data suggest that while synaptic density remains relatively homogenous across most sister branches, proximal compartments exhibit pronounced spatial fluctuations in spine density. This selective variance in excitatory, but not inhibitory, synapses implies a localized shift in the E/I balance across these specific dendritic subdomains.

Given that excitatory synapses are non-uniformly distributed, we next asked whether there is a finer patterning of synaptic connections within the different subdomains. Several studies have demonstrated that an increase in the volume of excitatory synapses is associated with synaptic strengthening^62–64^. Both structural (formation of elimination of spines) and functional (changes in spine head volume reflecting changes in synaptic weights) forms of synaptic plasticity seem to be influenced by and interact with neighboring pre-existing synapses, often resulting in the formation of the so-called synaptic clusters. Indeed, several studies have investigated how non-linear dendritic mechanisms promote plasticity in nearby excitatory synapses^24,57,60,61,65^. For this reason, we investigated the presence of structural clusters in the different dendritic subdomains. To do so, we first defined a putative structural synaptic cluster as a group of three or more large synapses (as defined previously) within a branch segment of approximately 10µm^36,66^. This approach is based on the assumption that heterosynaptic plasticity drives co-tuned nearby synapses to grow together and deviate from the average size^33,39,57,67–69^. We used a hierarchical clustering algorithm to identify these putative structural clusters (**Figure 2E**). Our analysis revealed a higher density of structural clusters in the terminal domains, a result that was deviated significantly from the density distribution obtained through domain-specific shuffling of synaptic distribution (**Figure 2F**). However, when comparing these observations with a subdomain-specific null distribution, the differences were no longer significant (**Figure 2F**). This finding suggests that the observed variations in cluster density between dendritic domains are primarily driven by differences in synaptic density within the respective subdomains.

Taken together, our results indicate that the spatial distribution of excitatory synapses in CA1 PNs is not random; rather, it shows a clear enrichment in terminal branches of basal and apical oblique dendrites, but not in apical tuft dendrites. Notably, this fine, domain-specific clustered organization was not found for inhibitory synapses, creating an asymmetric excitatory-inhibitory (E/I) balance.

### From Reconstruction of Synaptic Distribution to Modeling of Neuronal Functional Properties

To investigate how E and I synaptic distributions may influence the emergence of spatial tuning in CA1 pyramidal neurons, we constructed a biophysical model for each reconstructed neuronal morphology (**Figure 3A**) and simulated their activity during a virtual linear track exploration task (**Figure 3B**). The models were implemented using the NEURON simulation environment^70^, adhering to the channel distribution parameters previously established^26^ **(see Methods**).

**Figure 3:**
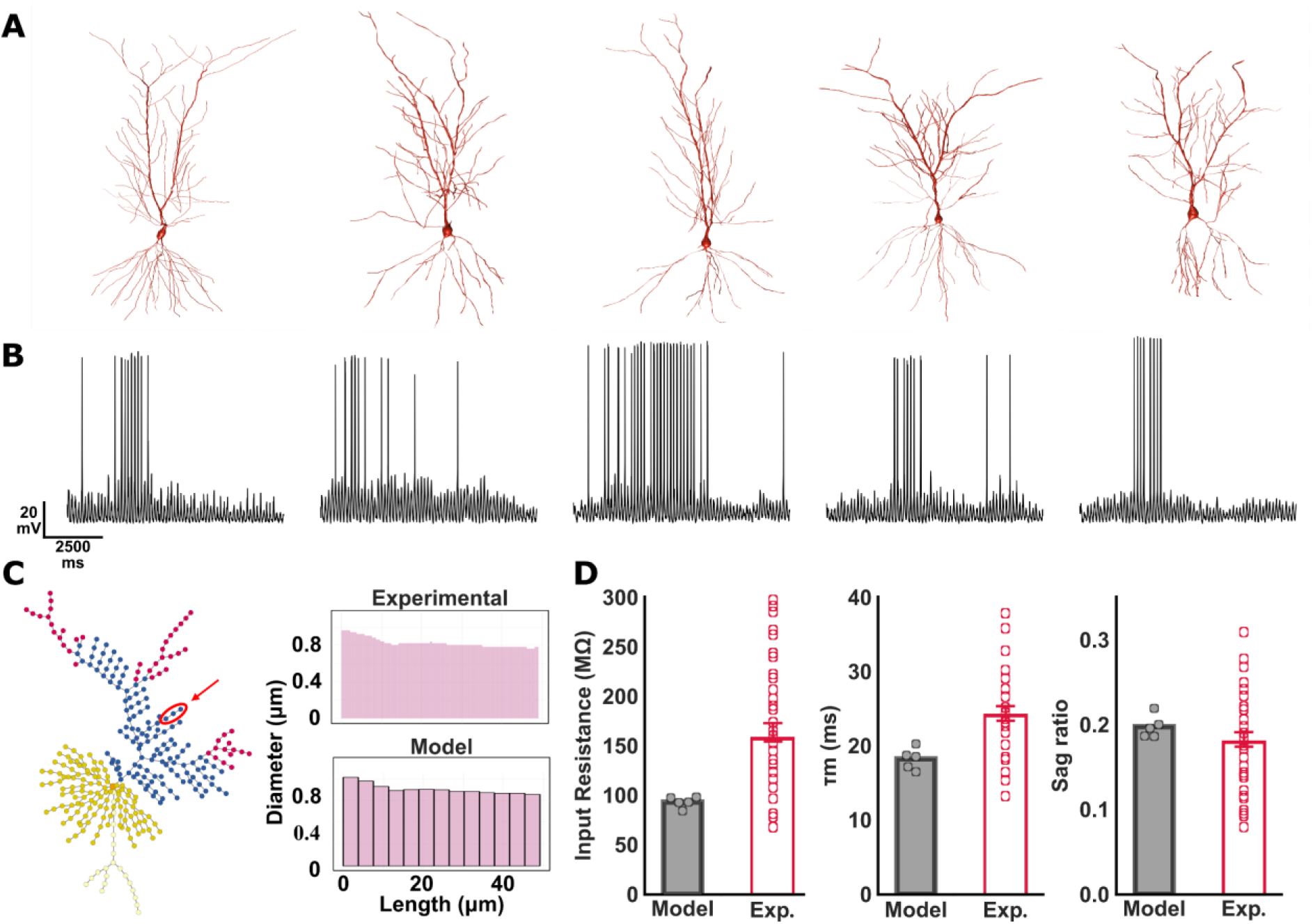
From morphologies to biophysical models. **A.** Reconstructions of dendritic morphology of the five complete CA1 pyramidal neurons used to generate the computational models. **B.** Somatic voltage traces produced by a place-tuned stimulation protocol used on the corresponding neuronal reconstruction. **C.** Left: schematic illustration of the cell’s segmentation used in the NEURON simulation environment (obtained using DendroTweaks^70^). Right: example of a branch structure from the data and its corresponding segmentation in the simulator. **D.** Passive membrane properties of CA1 PN models match experimental values. From left to right: input resistance, membrane time constant, and sag ratio. Experimental barplots redrawn from Masurkar et al., 2020^8^.

Given the high synaptic density in our reconstructed neurons, we employed a fine spatial discretization by increasing the number of compartments in the models. This approach achieved enhanced spatial resolution, allowing us to capture the detailed biophysical interactions between adjacent synapses (**Figure 3C**). To further ensure the accuracy and biological relevance of our models, we validated both their passive and active properties against experimentally observed values. Passive properties, including input resistance, membrane time constant, sag ratio, and dendritic attenuation, were all within the previously reported physiological ranges^8,71,72^ (**Figure 3D-S2A; see Methods**). Similarly, the active properties, such as somatic and dendritic spiking activity, Frequency-Current (F-I) curve, and dendritic nonlinearities, were validated using previously published protocols^8,24^ (**Figure S2B-E; see Methods**).

The excitatory synaptic weights were calibrated based on experimental findings to ensure physiological relevance^73^. Specifically, we implemented the principle of synaptic democracy, which posits that synaptic inputs across the dendritic tree contribute equally to somatic depolarization, regardless of their location^74^. To achieve this, proximal synapses were parametrized to elicit a 0.2 mV depolarization at the soma^20^. For Apical and Basal synapses, the α-amino-3-hydroxy-5-methyl-4-isoxazolepropionic acid (*AMPA*) and N-methyl-D-aspartate (*NMDA*) receptor conductances were progressively increased as a function of the distance from the soma^73,75,76^. This adjustment ensured that while distal synapses generated larger dendritic excitatory postsynaptic potentials (EPSPs), the resulting somatic depolarization remained constant at 0.2 mV (**Figure S2F**; **see Methods**). Moreover, considering the large variability of spine sizes observed in our analysis, we decided to account for it in the calculation of synaptic strength by linearly varying the validated weight according to the spine head volume^77^ (**see Methods**).

Inhibitory synapses were validated differently, based on their presynaptic sources, to ensure an accurate representation of their functional characteristics. For this study, inhibitory synapses were assigned to six distinct interneuron types, neurogliaform (NGF), oriens-lacunosum moleculare (OLM), parvalbumin-expressing basket (PVbasket), apical dendrite targeting cholecystokinin-expressing (CCK), Bistratified, and Ivy cells, based on their known post-synaptic target distributions across the dendritic tree^78–80^. The activity of all interneuron models was modulated by oscillatory rhythms, as suggested by experimental data^79^(**Figure 5C**). Synaptic weights, as well as rise and decay times, were calibrated by replicating experimental protocols reported in the literature^81–85^ (**Figure S2G**). Similar to their excitatory counterparts, inhibitory synapses were also scaled linearly with measured synaptic volume. These rigorous validations enhanced the reliability of the models and ensured they accurately reflected the physiological characteristics of CA1 PNs.

### Synaptic clustering amplifies dendritic gain and sustains calcium influx under noisy input regimes

Previous studies have suggested that synaptic clustering greatly influences the probability of dendritic spiking, especially under conditions of coincident input^24,26,86^, and leads to strong neuronal output, suggesting a mechanism for the amplification of salient input signals. Thus, to assess the functional impact of the observed, non-uniform, synaptic distribution on neuronal output, we simulated the activity of our biophysically detailed CA1 PN models using noisy inputs. We systematically stimulated an increasing number of excitatory synapses with independent, Poisson-distributed inputs, each with an average firing rate of 5 Hz (**Figure 4B**), while comparing two specific spatial distributions: a Clustered configuration, based on our morphological data, and a Shuffled one (**Figure 4A**). In the Shuffled configuration, the spatial location of activated synapses was randomized across the dendritic arbor, while preserving identical spine volumes and presynaptic activation patterns to isolate the effect of spatial organization. Analysis of the resulting topologies, via the use of hierarchical clustering (as in **Figure 2E**), confirmed that the Clustered configuration maintained a significantly higher density of spatial clusters compared to the Shuffled configuration across the full range of synaptic activation levels (**Figure 4C**). In line with previous studies^24,26,29,36,86–91^, our simulations revealed that the spatial clustering of inputs serves as a powerful amplification mechanism. Specifically, for any given number of activated synapses, the Clustered configuration consistently drove the neuron to significantly higher somatic firing frequencies compared to the Shuffled one (**Figure 4D**). This enhanced somatic output was caused by the elevated dendritic activity (**Figure 4E**), confirming that clustered inputs more effectively engage local dendritic integration mechanisms. However, the relative advantage conferred by clustering did not scale linearly with the number of activated synapses. Instead, Clustering offered a significant benefit for a relatively narrow range of input densities (3-9%), peaking at 6%. As the number of activated synapses increased beyond this range, the performance of the Shuffled configuration progressively approached that of the Clustered configuration, ultimately yielding comparable dendritic firing rates (**Figure 4D-E**). This convergence is likely due to the gradual increase in the number of activated synapses, which can generate enough depolarization to drive dendritic nonlinearities irrespective of their spatial distribution. The latter is also evidenced by the normalized difference in sodium charge (*Na^+^*) and NMDA spike counts, which drop towards zero in the high-input regime (**Figure 4H**). Interestingly, certain dendritic dynamics remained unmatched even when somatic and dendritic firing rates (**Figure 4D-E**) and fast sodium-mediated events (**Figure 4G**) became comparable between configurations. Specifically, the Clustered configuration maintained a distinctly higher level of voltage-gated *Ca^2+^*influx (**Figure 4F**) and total integrated NMDA charge (**Figure 4I**). This difference plateaued rather than diminishing, suggesting a fundamental decoupling of somatic spiking and dendritic biochemical signaling at high input rates. These results indicate that while distributed inputs can drive similar dendritic and somatic spiking if sufficiently numerous, only the Clustered configuration ensures a sustained, elevated state of dendritic *Ca^2+^*signaling. Given the essential role of *Ca^2+^* signaling in long-term potentiation (LTP) and synaptic strength maintenance^92–97^, this suggests that the primary computational benefit of the observed synaptic clustering may be to lower the threshold for plasticity-related calcium events, thereby facilitating the stable encoding of place fields even under noisy or sparse input conditions. This prediction adds an important functional role to synaptic clustering that goes beyond signal amplification and opens new avenues for experimental investigations.

**Figure 4:**
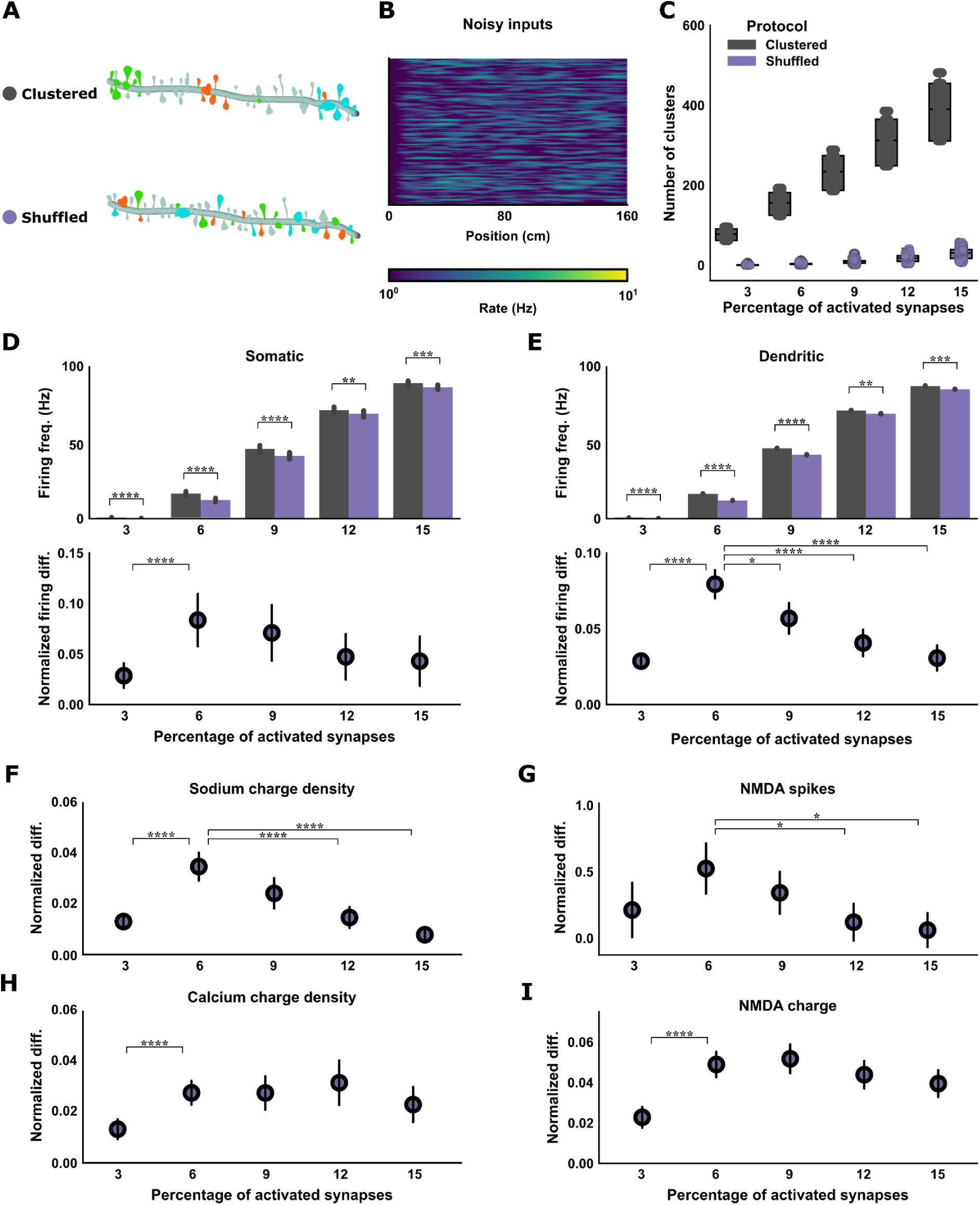
Spatial Synaptic Clustering Amplifies Somatic Output and Sustains Dendritic Signaling. **A.** Schematic representation of the Clustered and Shuffled input configurations. **B.** Representation of the stimulation inputs: 5 Hz Poisson-distributed inputs were delivered to an increasing number of synapses. **C.** Quantification of synaptic clustering confirms that the Clustered configuration retains a higher number of clusters than the Shuffled configuration as the percentage of activated synapses increases. Synapses were activated across a range of 3% to 13% of the total synaptic population (in 3% increments), corresponding to an average increase of ∼350 synapses per step. **D-E.** Top: Somatic (D) and Dendritic (E) firing frequency in the Clustered and Shuffled configurations. Spatial clustering consistently drives higher somatic and dendritic output. Bottom: normalized differences between Clustered and Shuffled configurations in Somatic (D) and Dendritic (E) firing rate. Clustering effect on firing rate is maximized around 6% of activated synapses (for all comparisons Wilcoxon signed-rank test was used). **F–I.** Decoupling of Spiking and Calcium Signaling: Normalized differences between Clustered and Shuffled configurations of fast sodium-mediated charge density (F) and ***NMDA*** spike counts (G) show convergence at high percentages of activated synapses. Conversely, voltage-gated *Ca^2+^* charge density (H) and total integrated NMDA charge (I) normalized differences remain significantly elevated, even after dendritic output in the two configurations converges. For all groups, Games-Howell post-hoc was used. Significance: ∗p < 0.05, ∗∗p < 0.005, ∗∗∗p < 0.001, and ∗∗∗∗p < 0.0001. Error-bands in D, E, F, G, H and I represent 95% confidence interval (CI) of the mean.

### Functional Role of Input Clustering in Place Cell Computation

After confirming that excitatory synaptic clustering activates dendritic integration mechanisms more efficiently and drives somatic firing more effectively than shuffled synaptic distribution, we investigated how the spatial organization of excitatory inputs influences place-tuning properties. Specifically, we examined how the clustering of excitatory synaptic inputs alters a neuron’s computational capabilities during a simulated spatial navigation task.

Specifically, we simulated the input (from EC and CA3) and output dynamics of our CA1 PN model under a virtual 160cm linear track exploration task. Briefly, EC inputs exhibited grid-like activity while CA3 inputs exhibited place-like activity or background noise, as dictated by experimental data^1,2,98–100^. Both excitatory and inhibitory inputs were theta-modulated, as depicted in **Figure 5** (see **Methods** for details).

**Figure 5:**
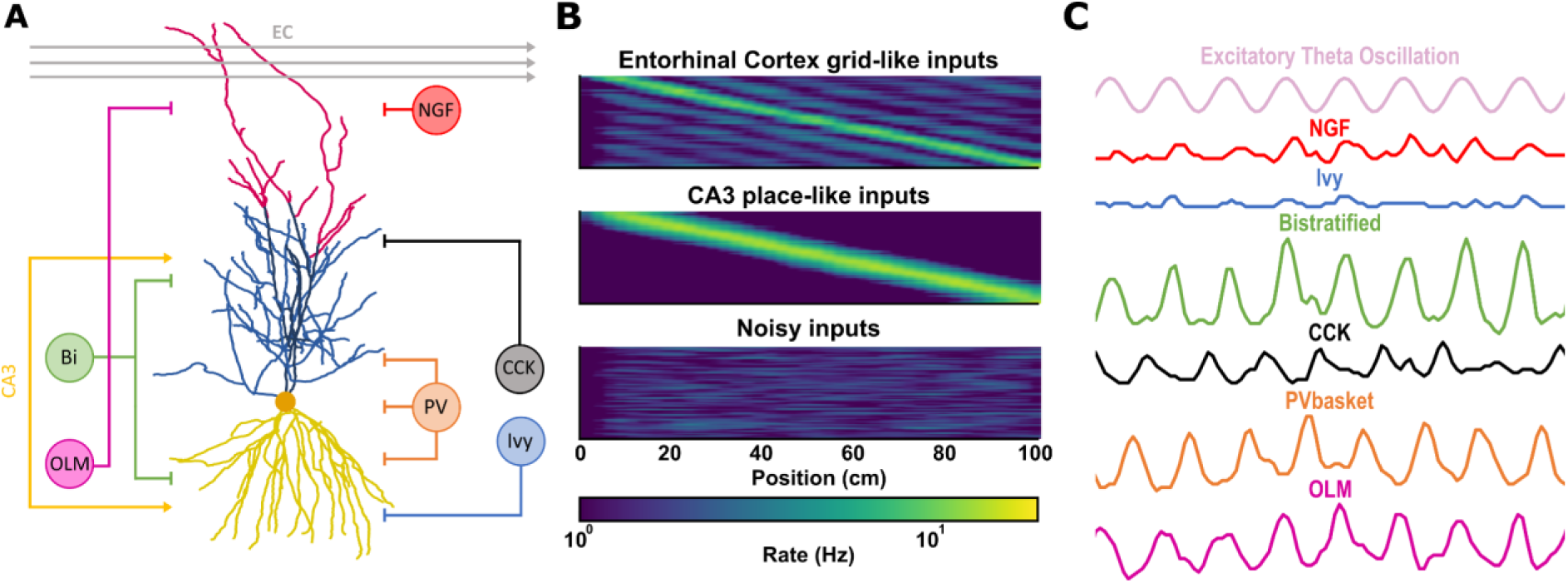
Patterns of Excitatory and Inhibitory Inputs to CA1 Pyramidal Cells. **A.** Schematic representation of the model neuron depicting excitatory and inhibitory input sources that are organized according to anatomical findings. **B.** Different types of excitatory inputs to the model. From top to bottom: Heatmap of EC grid-like, CA3 place-like, and random noise spike trains. Each row represents a different presynaptic input neuron. The colormap is the same across heatmaps, and brighter colors denote higher firing rates. **C.** Population activity of simulated interneuron spike trains compared to the Excitatory Theta Oscillation. Firing frequencies and theta phases are based on experimental findings.

The activity dynamics of excitatory inputs were also calibrated according to a recent study which characterized the spatiotemporal patterns of synaptic inputs to CA1 place cells during navigation on familiar linear tracks^41^. This study revealed that while synapses receive inputs across all track locations, the excitatory drive is significantly higher within the somatic place field than outside it. Furthermore, synapses co-tuned with the soma exhibit a greater tendency to form spatial clusters. Based on these findings, we generated 500 simulated connectivity patterns for each neuronal reconstruction designed to emulate the observed spatial organization. Specifically, each pattern included a subset of structural clusters preferentially targeted by place-tuned inputs. Inputs associated with a selected place field targeted a higher number of clustered synapses (designated as Functional Clusters), which determined the somatic place-tuning. The remaining synapses either remained silent, or received place-tuned or noisy inputs drawn from a uniform probability distribution (see **Methods** for Details). This distribution established our Clustered configuration (**Figure 6A**). Of note, prior studies used alternative designs, whereby infield synapses had a higher synaptic weight^101,102^. To evaluate the impact of clustering, we established two additional synaptic distributions. First, we generated a Shuffled distribution by randomly reassigning the locations of activated synapses within the same dendritic domain (**Figure 6A**, Shuffled). Initial simulations using this configuration revealed a significant reduction in overall neuronal activity (**Figure 6B and 6C**). To account for this loss of drive, we created a Shuffled - restored configuration. In this condition, we gradually increased the number of activated synapses by randomly sampling from the original input set, until we compensated for the observed activity drop. Restoration required approximately 13% more synapses compared to those activated in the Clustered configuration (**Figure 6G**).

**Figure 6:**
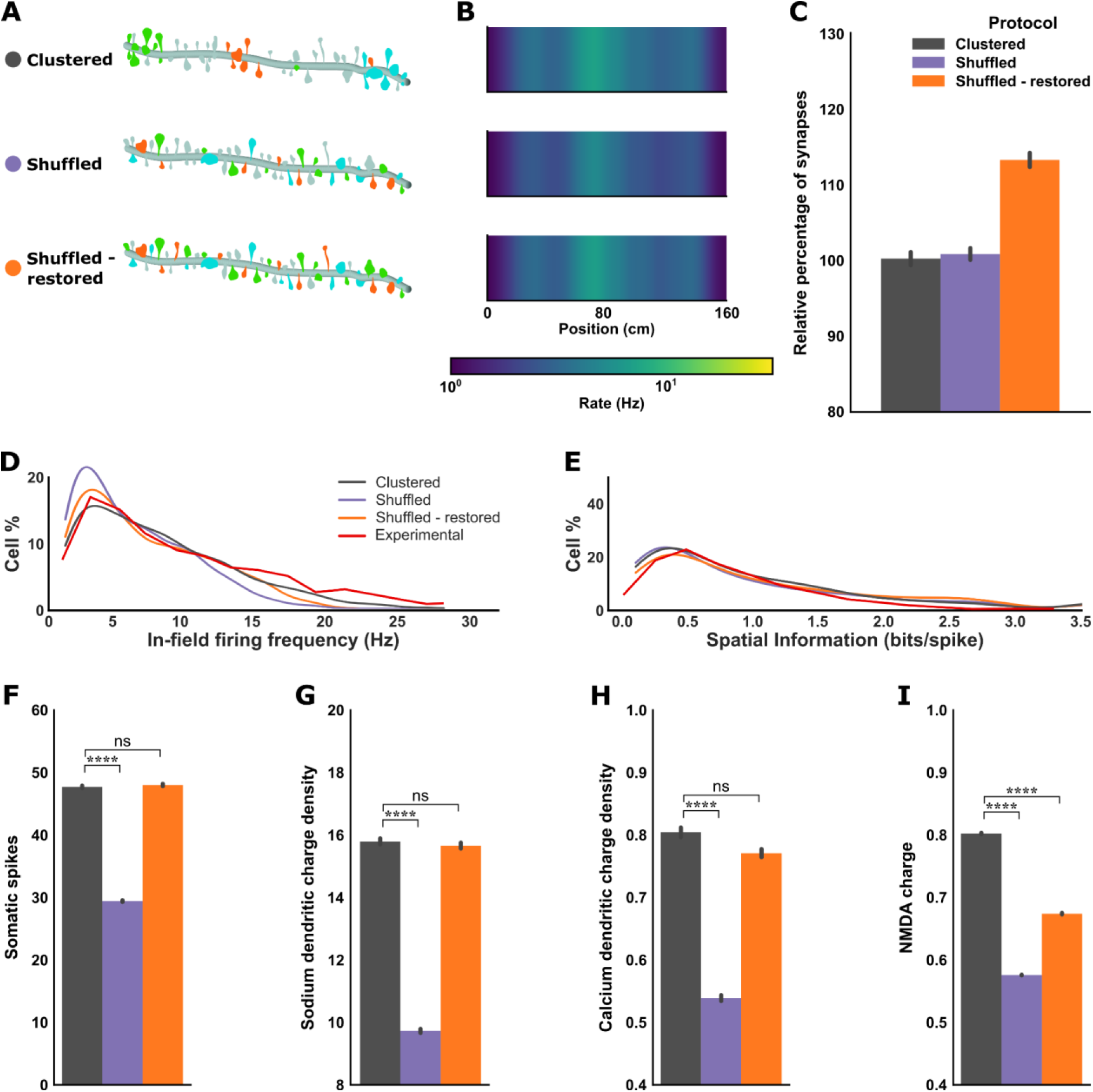
Effect of Synapse Clustering on Spatial Tuning. **A.** Schematic representation of three different synaptic configurations. Colors represent the tuning of synaptic inputs. Clustered: Synapses within pre-identified structural clusters are more likely to receive co-tuned inputs. Shuffled: activated synapses are randomly distributed within the same domains while maintaining the same inputs. Shuffled - restored: Same as in Shuffled but for a larger number of synapses to compensate for the reduced somatic activity. **B.** Activity heatmaps showing place-tuning of a population of CA1 PN models stimulated under the different synaptic configurations. **C.** Percentage of activated synapses for each synaptic configuration, relative to the Clustered distribution (Average for Clustered, Shuffled = 100%; average for Shuffled - restored = 113%). **D.** Distribution of in-field firing frequencies. Both the Clustered and the Shuffled - restored configurations qualitatively replicate the experimentally observed spatial firing patterns of CA1 PNs as reported by Mizuseki et al. (2012)^103^. The firing frequency distribution of the Shuffled configuration is skewed towards lower values. **E.** Distribution of spatial information content (measured in bits per spike). All configurations align qualitatively with the experimentally observed spatial firing patterns described by Mizuseki et al. (2012)^103^. **F.** Average somatic spike number under different synaptic configurations. Input shuffling significantly decreases the firing rate of the output place fields (non-normally distributed data: Kruskal-Wallis test between ‘Clustered’, ‘Shuffled’ and ‘Shuffled - restored’: H(2) = 176.64, p<10^−30^). **G-I.** Average dendritic and synaptic ionic influx under the different synaptic configurations: charge density for *Na^+^* (**G**) and *Ca^2+^* (**H**) and integrated charge for *NMDA* (**I**). While a significant effect was found for all ionic charges in the ‘Shuffled’ configuration, only the integrated *NMDA* charge was decreased in ‘Shuffled - restored’. (non-normally distributed data: Kruskal-Wallis test between ‘Clustered’, ‘Shuffled’ and ‘Shuffled - restored’: Na^+^ charge - H(2) = 254.66, p<10^−50^; Ca^2+^ charge - H(2) = 287.47, p<10^−60^; Na charge - H(2) = 451.01, p<10^−90^). For all groups, pairwise Mann-Whitney U tests with Benjamini-Hochberg (FDR) correction were used. Significance: ∗p < 0.05, ∗∗p < 0.005, ∗∗∗p < 0.001, and ∗∗∗∗p < 0.0001. Error-bands in C, F, and G represent 95% confidence interval (CI) of the mean.

To compare our models under these configurations, we used specific place field metrics, including in-field and out-field firing frequencies, field size, and spatial information (*SI*). To generate multiple examples of spatially-tuned neurons, we simulated different connectivity patterns and treated these simulations as a population of neurons, facilitating the comparison with experimental data^103^.

Our results indicated that all three configurations were qualitatively aligned with the SI distribution reported by Mizuseki et al.^103^ (**Figure 6D, 6E**). However, only the Clustered and Shuffled - restored configurations successfully matched the experimental in-field firing frequency. This suggests that while shuffling the synaptic positions reduces the firing rate, it does not inherently degrade the spatial information of the somatic output of our model cells. Finally, we analyzed the biophysical underpinnings of these differences by quantifying the dendritic *Na^+^* (**Figure 6G)**, *Ca^2+^* (**Figure 6H)**, and synaptic *NMDA* receptors (**Figure 6I)** ionic charge influx during the simulations. Consistent with the previous noisy stimulation experiments, the Shuffled configuration exhibited a sharp drop in all ionic charges. Notably, while the Shuffled - restored configuration succeeded in restoring *Na^+^* and *Ca^2+^* charge influx, it failed to recover the *NMDA* receptors levels (**Figure 6I**).

Overall, the above findings can be distilled in the following take-home messages: First, it appears that both spatially clustered and distributed synaptic configurations can generate high quality place fields with characteristics similar to those reported experimentally. However, a distributed configuration requires a much larger number of activated inputs to produce similar-quality place fields (**Figure 6C, D, E**). Second, the total synaptic drive, which can result either from few-and-clustered or from numerous-and-dispersed excitatory synapses, seems to be the key driver of dendritic and neuronal excitability. When the total synaptic drive is matched, both clustered and distributed synaptic configurations produce similar somatic outputs and dendritic excitability levels (**Figure 6F, G, I**). Third, synaptic clustering is a critical determinant of *NMDA* receptor activation and its associated *Ca^2+^* influx. Despite a match in total synaptic drive, the Clustered configuration drives a much higher *NMDA* receptor activation than the distributed ones (**Figure 6I)**.

### Dendritic subdomain-specific strategies differentially shape spatial tuning

While global synaptic clustering alters the place-specific properties of our model CA1 PNs, its role in domain-specific dendritic integration remains unclear. To elucidate the rules governing this phenomenon across our three distinct input distribution regimes, we employed a Random Forest (RF) multi-output regression analysis (see **Methods** for details) to predict somatic firing output (**Figure 7A, B**). Briefly, the RF model took as input domain-specific synaptic features of each place-tuned PN model (i.e., Interspine interval std, Number of clusters, Size of clusters, Path distance of clusters, Volume of clusters, Average synaptic volume) and used them to predict somatic output. Prediction accuracy was quantified using the Mean Squared Error (*MSE*) between the predicted and actual spatial information, in-field and out-of-field firing rates, and field size for each model neuron. The RF method achieved high predictive accuracy across all configurations (**Figure 7C**), allowing for a direct comparison of the feature importance rankings across the three synaptic distributions. This metric quantifies the degree to which each input variable reduces the model’s prediction error, effectively ranking the synaptic features by their functional impact on the somatic output. Our results reveal an important reorganization of the features driving somatic output among the three configurations (**Figure 7D**).

**Figure 7:**
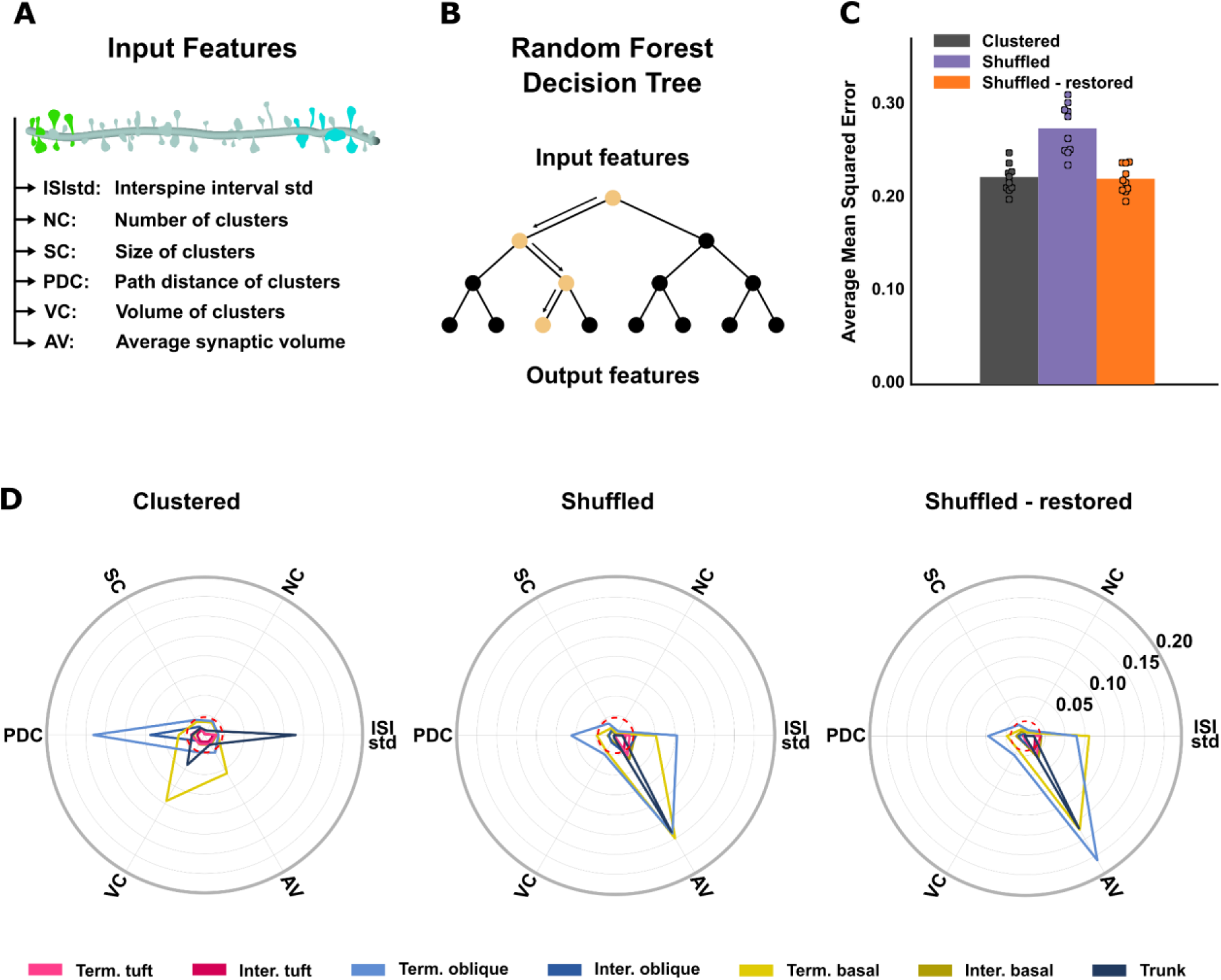
Synaptic features to Predict Spatial Tuning Quality Show Subdomain-specific Integration Strategies. ***A.*** Synaptic features used as input to the multivariate random forest model to predict spatial tuning quality. The latter is assessed by the spatial information, out-field firing frequency, In-field firing frequency and field-size metrics. For each dendritic domain (Basal, Apical, and Tuft), synaptic features include: intermediate to terminal ratio, Interspine Interval std, Number of clusters, Size of clusters, Path distance of clusters, Percentage of in-field branches, Average synaptic volume. **B.** Schematic representation of the random forest methodology. The model combines predictions from multiple decision trees, each trained on subsets of the data (2000 simulations used for training, 500 used for testing), to enhance robustness and accuracy in predicting model outputs. **C.** Average Mean Squared Error (*MSE*) of the output predictions on the testing set, providing a quantitative measure of the model’s prediction accuracy. The values were obtained by averaging the MSE of the four output parameters defining the spatial tuning quality. **D**. Feature importance in predicting simulated spatial tuning quality for the Tuft, Apical, and Basal dendritic domains of each synaptic configuration. The dashed line indicates the chance level of feature importance (∼0.023).

Specifically, in the Clustered configuration, the neuron does not employ a monolithic integrative strategy, but rather demonstrates a distinct compartmentalization of computational roles across dendritic subdomains. Within the basal compartment, predictive performance was primarily driven by synaptic strength metrics, specifically the average synaptic volume (*AV*) and the volume of clusters (*VC*). This suggests that the basal dendrites operate largely as a source of additive drive, where the total synaptic weight is the primary determinant of somatic output, regardless of the precise spatial arrangement within the branch. This finding aligns with models proposing that the basal tree scales the final somatic response through cumulative depolarization^88,104^. In contrast, both terminal and intermediate oblique branches heavily relied on the path distance of clusters (*PDC*) measured relative to the origin of the subdomain. This suggests that oblique dendrites possess localized computational "hotspots," where the recruitment of supralinear clusters at specific distances from the branch origin dictates their impact on the soma. Furthermore, the trunk was mainly characterized by the importance of the interspine interval standard deviation (*ISI_std_*), indicating a reliance on the spatial distribution of co-tuned synapses. The indication of the domains as distinct, compartmentalized processing units, is a results that well aligns with the existing literature^105^.

The transition to the Shuffled and Shuffled-restored configurations forced collapse of this integrative hierarchy. As spatial clustering was disrupted in the Shuffled configuration, the domain-specific specializations observed in the clustered state converged into a uniform reliance on global synaptic weight. In the Shuffled-restored condition, the model still relied almost exclusively on *AV* across all subdomains. This indicates that while the neuron can maintain functional degeneracy by achieving the same firing rate through different biophysical means, it does so by sacrificing topological sensitivity. Without the structural prerequisite of clustering, the neuron reverts from a compartmentalized processor to a simple summation machine, where the nuances of cluster location and local dispersion no longer contribute to the spatial tuning of the output. Of note, both Shuffled configurations also rely on *ISI_std_*. While the contribution of this feature is smaller, it suggests that the cell strives to maximize the benefits of synaptic clustering even when it occurs by random chance. Finally, across all tested synaptic configurations, features associated with the apical tuft demonstrated negligible importance in shaping somatic output (**Figure 7D**). While this consistent lack of influence could be viewed as a byproduct of specific spatial constraints, it more likely points toward a fundamental functional specialization within the CA1 architecture. These results suggest that while the tuft is essential for top-down modulation and the induction of plasticity, it may not contribute directly to the real-time execution of spatial tuning during the exploration of familiar environments^106–108^. Instead, spatial representation appears to be dependent on both the strength-dependent drive of the basal tree and the location-dependent non-linearities of the apical obliques.

## Discussion

In this study, we combined high-resolution spatial mapping of excitatory and inhibitory synaptic distributions with biophysical computational modeling to investigate how the spatial distribution of excitatory synapses influences the integrative properties of spatially-tuned CA1 PNs. Although the structural and electrophysiological features of these neurons have been well-documented *ex vivo*^8–10^, linking detailed spatial distributions of both E and I synapses along the entire dendritic tree to their functional properties has remained poorly explored.

Here, we mapped the complete spatial distribution of excitatory (E) and inhibitory (I) synapses throughout the entire dendritic tree of individual mouse CA1 pyramidal neurons of adult mice *in vivo*. We found that excitatory synapses were non-uniformly distributed, with higher spine density and significant clustering on terminal apical and basal dendrites, whereas inhibitory synapses were more uniformly arranged. With our data, however, we were not able to identify branch-specific distribution patterns as, for instance, the two-stage integration model described in Katz et al., (2009)^22^. Biophysically realistic models, validated by electrophysiological data, showed that clustered excitatory inputs produce higher firing rates and stronger place-tuned responses than shuffled inputs, especially when fewer synapses are active. Disrupting clustering markedly reduced somatic and dendritic activity, requiring ∼13% more active synapses to restore place-field responses. Importantly, while excitability can be fully restored, dendritic calcium dynamics cannot, reinforcing the notion that clustering plays a key role in calcium dependent plasticity processes. The models further predict that synaptic organization determines integration strategy: clustered inputs preferentially leverage apical dendrites, while uniformly distributed inputs rely on summation across basal dendrites. Overall, clustered synaptic organization enables more efficient and robust place-field formation with fewer synaptic resources, highlighting the critical role of synaptic spatial arrangement in CA1 neuronal computation.

### Detailed morphological and topological mapping allows in-depth understanding of synaptic architecture in CA1 pyramidal neurons

While previous studies established the general organizational principles of CA1 pyramidal neuron inputs, they relied on representative, local, sampling to estimate total synapse numbers and distribution. Megías et al. (2001)^9^ combined 3D light-microscopic reconstructions of complete dendritic arbors with serial electron microscopy of selected segments, calculating total counts by extrapolating observed local densities across the total length of specific dendritic subclasses. More recently, Bloss et al. (2016)^109^ utilized large-volume array tomography to map synaptic connectivity with high precision across more than 350 individual branches; however, their neuron-wide estimates were still derived by projecting these branch-level findings onto reconstructed dendritic domains. In contrast, our approach utilizes high-resolution confocal imaging and automated detection to provide a complete, neuron-wide, mapping of every excitatory (dendritic spines) and inhibitory synapse across the entire dendritic tree of individual CA1 pyramidal cells. This methodology, mirroring the recent work in mouse layer 2/3 cortical neurons^44^, moves beyond statistical extrapolation to capture the actual spatial distribution and potential local correlations of every synapse on a single cell. This level of detail is critical for identifying organizational features that may be masked by the averaging inherent in localized sampling and extrapolation methods.

Moreover, our computational approaches allowed us to further investigate sub-branch functional structures, such as excitatory synaptic clusters, in a comprehensive manner. Synaptic clustering has traditionally been studied through longitudinal *in vivo* imaging of dendritic spine dynamics, identifying clusters by the emergence of new spines during learning and behavior^33,35,42,110,111^. While these studies provide a vital link between synaptic plasticity and functional outputs, they are inherently limited by sparse sampling, as optical constraints typically restrict visualization to a small fraction of the total dendritic arbor.

Although our dataset offers a static snapshot rather than a temporal record of spine formation, we used synaptic volume (dendritic spine heads) as a proxy for structural correlation of synaptic weight. Specifically, we defined putative structural clusters as groups of three or more synapses with volumes exceeding the 80th percentile within a 10 µm dendritic segment. This criterion is based on the premise that heterosynaptic plasticity drives the co-localization and synergistic growth of co-tuned neighboring synapses^112^. These connections, where a single presynaptic axon forms multiple spatially clustered contacts on the same dendrite, exhibit significantly greater synaptic strength than single-axon connections, including increased spine volumes, larger PSDs, and higher release probabilities. Furthermore, the spines within these clusters possess highly correlated morphologies, suggesting that shared pre- and postsynaptic activity exerts a uniform influence on their structure. By specifically identifying clusters of large-volume synapses, our analysis specifically focuses on structural motifs that likely represent these potent, co-active functional inputs.

Our finding that terminal dendrites exhibit a higher spine density compared to intermediate dendrites aligns with prior work^9,109^ and results in higher probability of finding spine clusters in terminal as opposed to intermediate dendritic compartments. This disparity raises important questions about the mechanisms that enable the preferential formation of spines toward terminal regions. We propose that the smaller diameter and the sealed ends of terminal dendrites lead to elevated depolarization, thereby increasing the likelihood of synaptic plasticity events and subsequent spine formation, a specialization that may enhance the neuron’s ability to encode spatial information^10,113,114^. This hypothesis is supported by previous modelling and experimental work, whereby activation of inputs in terminal dendrites facilitates the induction of NMDA spikes, resulting in persistent firing in L5 prefrontal cortical pyramidal neuron models^115^. Similarly, in L2/3 pyramidal neurons of the primary visual cortex, NMDA spikes in the tuft dendrites promote orientation tuning^116^. Models predict that such NMDA spikes are more effectively induced by co-activated spines that are organized in clusters^86^.

### Spine clustering as an efficient mechanism for the amplification of salient signals

Our biophysical modelling approach generated several predictions: first, we predict that inputs targeting clustered synapses of physiological size and localization can more efficiently drive somatic activity compared to a randomized case. This finding is due to enhanced signal propagation, increased signal-to-noise and reduced attenuation that is achieved via the generation of dendritic non-linear events. The latter are more efficiently induced when spines are activated in clusters, due to the spatio-temporal summation of associated depolarizing signals^43,117,118^. The amplifying effect of synapse clustering reaffirms well established principles of supralinear dendritic integration observed both in experimental^24,36,87–90^ and computational studies^10,26,30,86,91^. However, unlike previous work^86^, we find that relatively small clusters of 4-8 synapses suffice to enhance somatic output (**Figure S4**). This is likely because prior work examined functional clustering of synapses with uniform strengths^86^, whereas we adhere to the physiological size and localization distributions of both excitatory and inhibitory synapses.

Moreover, this amplification effect is maximized when the percentage of activated synapses is relatively low (∼6%, see **Figure 4**). When a large enough pool of excitatory synapses is activated, irrespective of their spatial arrangement, the resulting dendritic nonlinearities and somatic outputs can be very similar. Interestingly, this equivalence does not hold for all dendritic conductances. The voltage-gated *Ca^2+^* charge and the total integrated *NMDA* charge remain much higher under Clustered than Shuffled configurations, enabling a consistently higher *Ca^2+^* influx. This finding reveals a mechanistic explanation for the proposed prominent role of synapse clustering in various plasticity-related mechanisms^39,90,118,119^.

Of note, while achieving similar somatic outputs with distributed and clustered inputs is possible, it is much less efficient (see **Figure 6**). To achieve the same, spatially-tuned, somatic output with randomly distributed synapses, the cell requires approximately 13% more active contacts, increasing energy consumption and reducing available space for additional synapses^110,120^. This strategy aligns with principles of economical wiring and metabolic efficiency, suggesting that neurons may have evolved to optimize both their structural and functional features in order to better solve complex cognitive tasks^121^.

While our results seem to conflict with the work of Basak and Narayanan (2018)^122^, in which they observed sharper tuning with dispersed rather than clustered synapses, we argue that this is likely due to the use of a very different clustering protocol. In their study, clustering was defined as confining the entire synaptic load to just one or two oblique dendrites, a hypothesis that seems at odds with experimental data. Intense compartmentalization can lead to depolarization-induced block, where sustained voltage saturation prevents the precise firing required for sharp tuning^123^. In contrast, our approach is closer -albeit not identical- to what they call a ‘dispersed’ model. Their most effective tuning was achieved when synapses were spread across several branches to allow for the initiation and propagation of dendritic spikes without triggering a branch-wide block.

### Dendritic domain-specific strategies for spatial tuning

Our models demonstrate that the quality of spatial tuning is dependent on the interplay between synaptic topology and domain-specific dendritic biophysics. In the Clustered configuration, the neuron leverages a highly diversified strategy where different subdomains prioritize distinct features to shape somatic output. For apical oblique dendrites, the path distance of clusters (*PDC*) relative to the branch origin emerged as the primary predictor of tuning quality (**Figure 7**), suggesting that these branches operate as discrete functional subunits where the precise localization of supralinear inputs is important for somatic recruitment^14,24^. In contrast, the basal domain appears specialized for a more distributive role, where predictive power is driven by synaptic volume and cluster strength rather than precise spatial arrangement. This reinforces the view that basal dendrites provide a potent, additive depolarization that scales the somatic response, acting as a reliable gain-control mechanism for the nonlinearities generated within the apical tree^88,104^. Furthermore, the unique sensitivity of the trunk to the interspine interval standard deviation (*ISI_std_*) suggests a role for the local dispersion of inputs of this compartment in gating the integration of co-tuned signals before they reach the soma. Finally, the negligible influence of the apical tuft hints at a functional specialization, where the tuft may be reserved for top-down modulation and the gating of plasticity during environmental learning, rather than the real-time execution of spatial tuning^106–108^.

However, as synaptic topology transitions from Clustered to Shuffled, we observe a collapse of these specialized domain strategies. Our findings reveal that the disruption of spatial clustering forces the neuron to abandon its compartment-specific logic in favor of a uniform, summation-based strategy. Crucially, even in the Shuffled-restored condition, where somatic firing is recovered, the reliance on topological features dramatically reduces, replaced by a more general dependence on average synaptic volume (*AV*). This shift indicates that while CA1 neurons exhibit functional degeneracy^122^, this robustness comes at a cost. In the Shuffled configurations, neurons operate as a simple global integrator, losing the nuanced control provided by dendritic compartmentalization and spatial precision^23,124^. This suggests that clustering is not merely a way to increase firing rates, but a requirement for the subdomain-specific computations that define the CA1 pyramidal neuron as a sophisticated multi-compartmental processor^105^.

In sum, this work provides an integrated framework that combines detailed anatomical synaptic mapping and computational modeling to unravel how synaptic organization shapes the cellular computations underlying spatial navigation. We demonstrate that the neuron’s integrative state is not fixed but is highly contingent on the spatial architecture of its inputs. By linking subcellular morphological metrics to macroscopic computational outputs, this study advances our understanding of the fundamental structural principles governing neuronal processing in the hippocampus.

## Resource availability

### Lead contact

Further information and requests should be directed to and will be fulfilled by the lead contact, Panayiota Poirazi

### Materials availability

This study did not generate new, unique reagents.

### Data and code availability

The source code that generates all simulations and Figures, as well as the data that support this study will be freely available upon publication on GitHub.

## Acknowledgments

We thank all members of the PoiraziLab for their valuable feedback on the project. Moreover, we thank Dr. Hanchuan Peng, developer of Vaa3D, for developing the pipeline that allowed us to obtain the reconstructed morphologies. This project has received funding from the European Union’s Horizon 2020 research and innovation programme under the Marie Skłodowska-Curie grant agreement No.860949; Stavros Niarchos Foundation (SNF) and the Hellenic Foundation for Research and Innovation (H.F.R.I.) under the 5th Call of “Science and Society” Action Always strive for excellence – Theodoros Papazoglou” (Project Number: 28056); H.F.R.I call “Basic research Financing (Horizontal support of all Sciences)” under the National Recovery and Resilience Plan “Greece 2.0” funded by the European Union – NextGenerationEU (Project Number: 014941); COFLEX 14941 HFRI P.N. 80147/17.1.24; NIH grant No 1R01MH124867 to P.P.; NIH grants F31NS101820, F32MH125600, and K99DK142063 to D.M.I.; NIH-NINDS (R35 NS127232) (FP), and an award from the NOMIS Foundation (FP). A.L. is supported by National Institute of Mental Health (NIMH) R01MH124047 and R01MH124867; National Institute on Aging (NIA) RF1AG080818; National Institute of Neurological Disorders and Stroke (NINDS) Brain Initiative U01NS115530; NINDS R01NS121106, NINDS R01NS131728, and NINDS Brain Initiative R01NS133381

## Author contributions

P.P., A.L., F.P., S.C., D.M.I., and S.T. conceived the study. D.M.I. performed the in-utero electroporation, prepared the tissue, and acquired the confocal images. L.H. developed a script enabling image deconvolution of the raw PN images. Y.K. and D.M.I. manually edited semi-automated synaptic reconstructions and participated in data analysis.

S.T. refined the morphological traces, developed and validated the model, implemented the simulation code, and analyzed the data. S.C. and S.T. generated the input patterns. S.T. wrote the manuscript, with editing support from P.P., A.L., F.P., S.C., and D.M.I.

## Declaration of generative AI and AI-assisted technologies in the writing process

While preparing this work, the authors used large language models (ChatGPT-4o, Gemini 3) for spelling and grammar checks. The author(s) reviewed and edited the content as needed and take full responsibility for the final publication.

## Materials and Methods

### Mice

All animals were handled according to protocols approved by the Institutional Animal Care and Use Committee at Columbia University, New York. Postnatal day 98 C57BL/6 mice (strain code: 000664; Jackson Laboratory) were used for all experiments. Timed-pregnant female mice were maintained in a 12-hour light/dark cycle and obtained by overnight breeding with males of the same strain. For timed-pregnant mating, noon after mating is considered E0.5.

### Constructs

The tdTomato reporter was previously generated by the Polleux lab^44^. Briefly, the tdTomato insert was subcloned into the pAAV-Ef1a-DIO eNpHR 3.0-EYFP plasmid (Addgene 26966) between the AscI and NheI cloning sites. The inhibitory synapse reporter EGFP-Gephyrin (clone P1) was obtained from H. Cline (TSRI, La Jolla, USA) and subcloned into the pUBC-DIO Teal-Gephyrin plasmid (Addgene 73918) between the AscI and NheI cloning sites.

### *In utero* electroporation

*In utero* electroporation targeting the dorsal hippocampus was performed using a triple-electrode setup as previously described in E15.5 mouse embryos^13,125,126^. Endotoxin-free DNA was injected using a glass pipette into one ventricle of the mouse embryos. Electroporation was performed at E15.5 using a square wave electroporator (ECM 830, BTX) with two anodes (positively charged) placed laterally on either side of the head and one cathode (negatively charged) rostrally at a 0° angle to the horizontal plane. The electroporation settings were: 5 pulses of 40V for 50 ms with 500 ms intervals. Plasmids were used at the following concentrations: Flex-tdTomato reporter plasmid: 1 μg/μl; Flex-EGFP-Gephyrin 0.45 μg/μl; NLS-Cre recombinase: 150 pg/μl.

### Tissue preparation

Animals at the indicated age were anaesthetized with isoflurane before intracardiac perfusion with PBS and 4% PFA (Electron Microscopy Sciences). As previously described^44^, 130 μm coronal brain sections were obtained using a vibrating microtome (Leica VT1200S). Sections were mounted on slides and briefly dehydrated at room temperature to reduce section thickness before being coverslipped in Fluoromount-G (SouthernBiotech).

### Confocal imaging

Confocal imaging was performed as previously described^44^. Briefly, images of electroporated neurons in slices were acquired with a Nikon AXR confocal microscope equipped with a 100x, 1.49 NA Apochromat oil-immersion objective and controlled by NIS-Elements (Nikon Instruments Inc., Melville, NY). Image volumes were acquired using galvano scanning with a 1024x1024 scan region, corresponding to a lateral pixel size of 110 nm and a z-step size of 100 nm. To compensate for signal attenuation caused by tissue scattering and spherical aberration with increasing depth, laser power was linearly increased as a function of depth to normalize the mean fluorescent intensity of pixels from image planes throughout the stack^44^. Following acquisition, image stacks were deconvolved using the GPU-based Richardson-Lucy deconvolution algorithm within NIS-Elements. Deconvolution was performed with a theoretical point spread function based on imaging acquisition parameters and used 20 iterations. Individual z-stacks were subsequently stitched together using Fiji’s Grid/Collection stitching tool^109^ to generate a single three-dimensional volume representing the entire neuron. Dendritic spines and inhibitory synapses were quantified based on tdTomato fluorescence and EGFP-Gephyrin puncta fluorescence, respectively. All quantifications were performed in CA1 hippocampus in sections of comparable rostro-caudal position.

### Neuron trace reconstruction

Neuron traces, digital reconstructions of the dendritic tree, are an effective representation of neuronal topology and geometry. The traces are described using a tree graph and consist of 3-D point coordinates, diameters, and connectivity between points. This representation enables an extensive quantitative analysis of the geometrical organization of the neurons they represent including total length, branching angles, distribution statistics and cumulative distance from the soma^127^. In this paper, the initial reconstructions are obtained using the automatic tracing methods built into the open-source 3D visualization and analysis tool Vaa3D^47^. Then, experts manually proofread the traces and make adjustments with the built-in proof-editing tools.

### Synapse detection

Excitatory and inhibitory synapses were annotated with the Spine Detector and IS Detector toolkits within Vaa3D software as previously described^44^. Briefly, excitatory synapses are identified as dendritic spines segmented by the Spine Detector toolkit using the image and neuron trace as inputs^128,129^. Because the dendrite traces represent the dendrites with a series of overlapping nodes^130^, information about the volume and distance from the dendrite of each spine can be associated with its nearest node to assign a location within the spatial context of the dendritic arbor. Users have the ability to accept or reject potential spines and adjust their volume.

We labeled inhibitory synapses using the scaffolding protein Gephyrin tagged with EGFP as a marker^48^. Because these synapses can only occur on the dendrites or the spines of neurons of interest, IS Detector uses the image from the cell-fill channel containing the dendrites and the spines as a mask image to extract the relevant region for the inhibitory synaptic marker. Then, signal beyond user-input parameters for minimum/maximum voxel count and distance from the trace is excluded and potential inhibitory synapses from the resulting image are identified based on a user-input intensity threshold. Users have the ability to accept or reject potential inhibitory synapses, adjust their volume, and assign them as dendrite-targeting or spine-targeting.

### Dendritic diameter correction

To improve the accuracy of the morphological data and ensure the physiological plausibility of the neuronal reconstructions, a custom smoothing algorithm was applied to correct artifactual fluctuations in dendritic diameters (**Figure S1**). This algorithm utilized a weighted moving average filter to smooth out large changes in diameter while preserving the overall structural integrity of the neuron. The filter was designed to account for local variations in diameter by applying larger weight to nearby points, reducing the influence of outliers and providing a consistent tapering pattern along the dendrites.

Prior to smoothing algorithm, a minimum diameter threshold was applied to all diameters to address the occurrence of bottlenecks. These bottlenecks could significantly impact simulations by increasing the axial resistance and altering signal propagation. The minimum diameter threshold was domain-specific, reflecting the physiological constraints of different neuronal compartments.

### Synaptic distribution analysis

#### Synaptic null distribution: within-domain and within-subdomain randomization

For each reconstructed morphology, synapses were randomly reassigned within branches of the same domain or sub-domain, ensuring a uniform probability distribution. Specifically, we first extracted the nodes of interest from the reconstructed arbor trace and then reassigned the annotations describing spine features to new, randomly selected locations within the same domain. This process preserved the overall structure of the arbor while randomizing synaptic positions. We repeated this procedure 1,000 times to create the null distribution, conducting the necessary analysis on each randomized dataset.

#### Structural cluster analysis

Putative structural clusters were identified using a hierarchical clustering algorithm applied solely to large spines, i.e., those above the 80th percentile of the spine volume distribution. For each branch, we selected the large spines and clustered them based on their relative distance from the soma (i.e., path length). Specifically, we used the SciPy hierarchical clustering linkage with *complete* method to calculate the Euclidian distance. A threshold of 10 micrometers was then used to define clusters, after which we collected their characteristics (e.g., spine count, total volume). Since the clustering was performed separately for each branch, any potential clusters spanning branching points were excluded from the analysis.

### Computational Modeling

The CA1 PN models, implemented using the Python version of the NEURON simulation environment^70^, and were inspired by prior work by using the ionic mechanism and their distribution throughout the dendrosomatic axis^26^. The ionic channels, written using NMODL language, used for this project include: inward transient sodium current (I_Na_), outward delayed rectifier potassium current (I_Kdr_), A-type potassium current (I_A_), Sodium persistent current (I_Nap_), Hyperpolarization-activated current (I_h_), Slowly activating muscarinic potassium current (I_m_), Calcium current from T-type channel (I_CaT_), Calcium current from L-type channel (I_CaL_), Calcium current from R-type channel (I_CaR_), Slow calcium-dependent potassium current (I_sAHP_), Medium calcium-dependent potassium current (I_mAHP_). Minor modifications were implemented in the kinetics and distribution of some channels. To facilitate the rapid and reproducible generation of models, we developed a Python Class capable of processing neuronal morphologies in .swc file format and automatically incorporating the appropriate biophysical properties and channel distributions (**Supplementary Methods, Table S1**).

To ensure high spatial resolution in the simulations, we determined the number of segments (*n*_*seg*_) for each neuronal section using the lambda rule, which is based on the Alternating Current (AC) length constant (*λ*). A ten-fold higher frequency was used in the calculation to obtain a finer spatial discretization of each section. Specifically, we employed the following equation (Eq. 1):

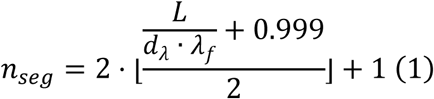

where *L* is the length of the section, *d*_*λ*_ the fraction of the AC length constant (typically 0.1), *λ*_*f*_*f* the AC length constant at the chosen frequency *f*, and ⌊⋅⌋ denotes a function that converts a specified value into an integer number. This approach ensured that all segments were small enough to disentangle inputs coming from most of nearby synapse couples, providing accurate spatial and biophysical representation of the model.

### Modeling Synapses

We incorporated both glutamatergic (excitatory) and GABAergic (inhibitory) synapses. All dendritic spines identified in the reconstructed neurons were assumed to be excitatory synapses containing both AMPA and NMDA receptors, while inhibitory synapses were modeled as GABA_A_-mediated inputs. Synaptic strength was adjusted based on the synapse size measured from the original reconstruction and, for a subset of excitatory synapses, by the distance from the soma. Specifically, for excitatory synapses, validated conductance values for both NMDAR and AMPAR components were first calculated. The AMPA conductance was then scaled by a volume-dependent scaling factor (Eq. 2). Both AMPA and NMDA components for synapses in Basal, Trunk and Oblique compartments were further adjusted by a distance-dependent scaling factor^73,75^. While these components have been previously characterized as a function of dendro-somatic distance in relation to the presence of perforated or non-perforated postsynaptic densities^131^, our fluorescence-based structural dataset lacked the resolution to distinguish between these synaptic subtypes. Consequently, we maintained a neutral biophysical assumption, scaling AMPA and NMDA components solely by measured spine volume and dendro-somatic distance. In contrast, inhibitory synapses were scaled solely by the volume-dependent factor (Eq. 3).

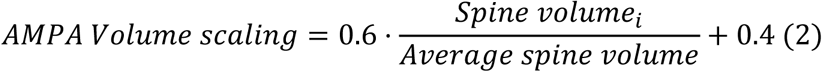

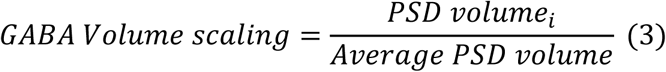

#### AMPA & GABA_A_ definition

AMPA and GABA_A_ synapses were simulated with a two-state kinetic model, defined by rise and decay times. The synaptic current is calculated according to the following equations (Eqs. 4, 5):

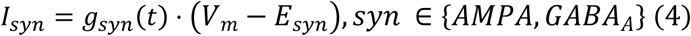

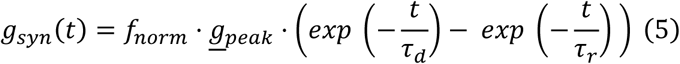

where *I*_*syn*_ is the synaptic current, *V*_*m*_ the membrane potential, *g*_*syn*_ the receptor conductance, *g*_*peak*_ the peak conductance and *E*_*syn*_ its reverse potential. *τ*_*r*_ and *τ*_*d*_ represent the rise and decay time constants, respectively. The factor *f*_*norm*_ normalizes the difference of exponentials so that the peak of *g*_*syn*_ equals <u>*g*</u>_*peak*_ and its calculated using the formula below (Eq. 6).

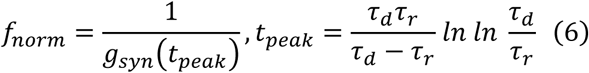

#### NMDA definition

The NMDA component of the excitatory synapses was also simulated as a two-state kinetic model but with an additional factor that accounts for the voltage dependent removal of the Magnesium blockage. The mechanism is described by the following equations (Eqs. 7, 8):

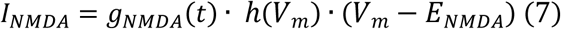

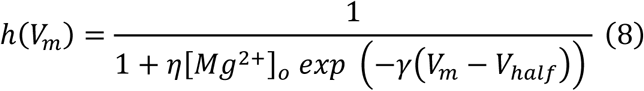

where *h*(⋅) is a sigmoid function, *η* denotes the Magnesium sensitivity of unblock, [*Mg*^2+^]_*o*_ the Magnesium extracellular concentration, *γ* the slope factor that determines how steep the transition is around the midpoint, and *V*_*half*_ the membrane potential of the midpoint of the sigmoid. *g*_*NMDA*_(*t*) is calculated with the same equations as the other synaptic currents (Eqs. 5, 6).

### Pyramidal neuron model validation

The parameters that defined the biophysical properties and channel distributions were carefully validated to match the cell’s behavior with experimental data. We ensured that using the same values for all morphologies would still adhere to the experimental values for each cell, while also introducing variability expected in a heterogeneous population of pyramidal neurons. We used hyperpolarizing step current sweeps to validate passive properties as the input resistance (*R*_*in*_), membrane time constant (*τ*_*m*_), sag ratio and the dendritic attenuation, following the protocols illustrated in **Magee (1998)**^8^**, Golding et al. (2005)**^72^ for the voltage attenuation and **Masurkar et al. (2020)**^8^ for *R*_*in*_, *τ*_*m*_ and sag ratio. We then used a depolarizing step current sweeps to validate the frequency-input (F-I) curve and the adaptation from **Masurkar et al. (2020)**^8^. For the synaptic strength validation, we followed the protocol used in **Magee and Cook, 2000**^73^, stimulating average size synapses at increasing distance from the soma and maintaining the 0.2 mV somatic depolarization observed experimentally.

### Inputs design

We modeled convergent EC inputs as multiple grid representations covering a virtual linear track^6^ and CA3 place-tuned inputs as Gaussian filtered Poisson spike trains parameterized by peak location, firing rate peak amplitude, and field width^102^. The two resulting input streams produced 40 groups of similarly tuned spike trains, with each group covering the entire track with centers placed every 4 cm (**Figure 5B**). Furthermore, independent Poisson-distributed inputs were introduced to the CA1 PNs to simulate background synaptic noise (**Figure 5B**).

#### Grid-like and Place-like inputs structure

In accordance with the formulation described in **Shuman et al. (2020)**^132^, entorhinal cortex layer III (ECIII) grid-like inputs were modeled as the summation of three sinusoidal wave components, following the approach of **Solstad et al. (2006)**^133^. Specifically, grid-cell activity was represented as the sum of three two-dimensional sinusoidal gratings, each defined by a distinct wave vector. The resulting grid-like function was computed as (Eq. 9):

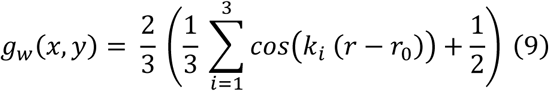

where the parameter set *w* = (*λ*, *θ*, *r*_0_) defines the grid spacing *λ*, orientation *θ*, and the spatial location of a theoretical place cell *r*_0_ = [*x*_0_, *y*_0_]^*T*^, while the variable *r* = [*x*, *y*]^*T*^ represents the current position, and *k* denotes the wave vectors arranged at angular intervals of 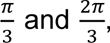 respectively (Eq. 10).

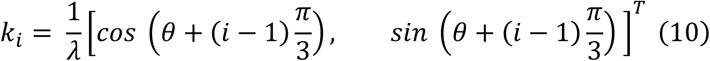

These grid-like functions were used to compute the probability of spike generation as a function of the animal’s position and the theoretical place fields. To incorporate theta rhythmic modulation, a sinusoidal filter was applied to these probabilities (Eq 11):

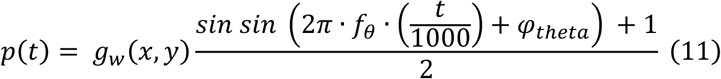

where *f*_*θ*_ = 8*Hz* and *ϕ*_*θ*_ = 0^*o*^ correspond to the theta-cycle frequency and phase, respectively, and *t* is expressed in milliseconds. A spike was generated for a given grid cell whenever *p*(*t*) exceeded a random value sampled from a normal distribution with mean *μ* = 0.7 and standard deviation *σ* = 0.05 or a random value from a uniform distribution used to introduce baseline noise. This process was repeated across all grid-like cells to construct the entire set of theoretical place fields.

To simulate CA3 place-cell activity, we modeled spike trains using a Poisson process gated by a spatiotemporal probability function. Unlike the EC-derived inputs, CA3 activity was generated using a Gaussian spatial tuning curve centered at the place field location and parametrized by firing rate peak amplitude and field width. For each synapse, an initial spike train was generated via a Poisson process (30–40 Hz), which was then filtered based on a probability *p*(*t*) determined by the product of the Gaussian spatial field and a sinusoidal theta-phase modulation function. As for EC inputs, spike was stochastically retained if *p*(*t*) exceeded a normally distributed threshold or if a random variable fell below the noise baseline. Finally, to ensure biological realism, we enforced a refractory period (*t*_*ref*_) to eliminate overlapping spikes and applied a 0.2 ms temporal jitter to the final spike times.

#### Noisy inputs

To account for non-place-tuned inputs, we generated a set of noisy inputs modeled as Poisson-distributed spike trains with low baseline firing rates. To introduce temporal fluctuations, these spike trains were modulated by applying a Gaussian filter at randomly selected time points, transiently increasing the firing rate. In our approach, the probability for a spike to occur was (Eq. 12):

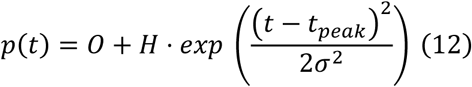

where *O* represents the baseline offset of the gaussian curve, *H* is the peak amplitude, *t* is the time at which the spike occurred and *t*_*peak*_ is a randomly chosen value corresponding to the mean value of the Gaussian filter. The standard deviation σ is also chosen randomly in a fixed window of values (400 ms, 720 ms) and represents the width of the window with slightly higher firing frequency.

This process ensured that noisy inputs exhibited sporadic increases of activity while maintaining an overall unstructured firing pattern, effectively capturing the background synaptic activity present in the hippocampal CA1 place cells. Similarly to place-tuned inputs, the noisy spike trains were further processed to account for synaptic failure.

#### Inhibitory inputs

Inhibitory inputs were modeled as theta-modulated, gamma-filtered Poisson-distributed spike trains, with firing properties tailored to different interneuron subtypes as described in **Bezaire et al., 2016**^79^. To generate these spike trains, we set baseline firing rates according to known physiological values for each interneuron type. These spike trains were then filtered using a theta-modulated function, ensuring that their activity exhibited phase locking to the theta cycle at subtype-specific phases (Eq. 11).

Additionally, a gamma-frequency filtering step was applied to refine the temporal structure of the remaining spikes. The level of randomness was controlled by introducing a noise parameter, which varied across interneuron types to reflect their intrinsic firing variability. Delays of 300 ms were incorporated to approximate transmission latencies. Specifically, inputs were tuned to replicate the activity of different interneurons neurogliaform (**NGF**) cells, oriens-lacunosum moleculare (**OLM**) cells, parvalbumin-expressing basket (**PVbasket**) cells, apical dendrite targeting cholecystokinin-expressing (**CCK**) cells, Bistratified cells, and Ivy cells with peak firing rates ranging from 2.8 Hz to 30 Hz, distinct theta-phase preferences, and varying degrees of noise, capturing the diversity of inhibitory influences in the hippocampal network (**Figure 5C**).

### Input distribution

Synaptic inputs were distributed in an anatomically accurate manner. For inhibitory synapses, their connection probabilities to each interneuron subtype were based on estimates from **Bezaire and Soltesz (2013)**^134^, with slight upward adjustments to account for missing interneuron types. While NGF and OLM inputs were uniformly distributed across tuft synapses, the distribution of other interneurons followed a nonlinear function of the distance from the soma. This distribution pattern was informed by anatomical observations while maintaining symmetry between the apical and basal dendritic domains. Specifically, PV basket synapses were concentrated near the soma, while CCK and Ivy cells targeted more distal synapses in the apical and basal domains, respectively. Bistratified cell synapses were predominantly located in the middle regions of both domains (**Figure 5A, S3**).

ECIII grid-like inputs were directed to synapses on the tuft dendrites in the stratum lacunosum-moleculare, whereas CA3 place-like inputs primarily targeted synapses in the trunk, oblique, and basal dendritic domains, spanning the stratum oriens and stratum radiatum. To distribute afferent inputs across the dendritic tree, synapses were categorized into place-tuned and noise-driven populations, each consisting of a mixture of clustered and randomly distributed synapses. For place-tuned inputs, synapses were sampled from the available pool based on predefined fractions for clustered 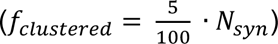 and random 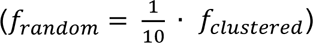 configurations. These fractions were chosen to replicate the distribution of place field properties observed at the population level by Mizuseki et al. (2012)^103^. We implemented a domain-specific sampling bias where synapses located on the apical tuft were sampled at half the frequency of those in other dendritic domains. Each place-tuned synapse was assigned to a specific place field (1 … *N*) by sampling from a probability distribution centered on a "chosen" place field. This distribution was defined by a Gaussian function (*sigma* = 2) shifted by a constant baseline (+0.6) to ensure an in-field to out-field firing ratio of 1.6, consistent with experimental observations^41^.

The remaining synaptic pool was sampled to populate the noise-driven input stream. Noise synapses were selected from the clusters and random nodes not previously assigned to place-tuned inputs, following the same fractional distribution logic (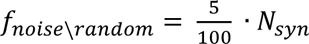 and 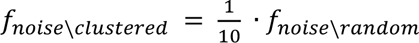). This approach ensured a non-overlapping assignment of inputs while maintaining the structural heterogeneity of the dendritic tree. For each reconstructed cell we generated 500 different configurations to ensure output variability.

### Synapse Shuffling

To isolate the functional contribution of synaptic clustering to neuronal output, we implemented two shuffling protocols that redistributed synaptic inputs while preserving domain-specific densities and input tuning. In the Shuffled condition, 500 unique connectivity matrices were generated for each reconstructed cell. Synapses were stochastically reallocated to new locations within their dendritic domains (Tuft, Apical Trunk, Apical Oblique, or Basal), thereby disrupting spatial clusters while maintaining the total number of inputs per domain.

The Shuffled - restored condition followed the same randomization procedure but included a ∼13% increase in the total number of activated synapses. This specific scaling factor was determined via a grid search to ensure that the resulting place-field properties at the population level remained consistent with the experimental distributions reported by Mizuseki et al. (2012)^103^.

### Place cell quantification

To quantify the spatial tuning properties of the simulated neurons, we computed the firing rate maps for each cell across the linear track, and used them to calculate the various place field features. The Field Size identifies the spatial extent of contiguous bins around the peak firing location where the firing rate remains above a given relative threshold. Once the Place Field width was characterized, we determined the mean firing rate within and outside the identified field region, named In-field and Out-field Firing Frequency, respectively. In addition, spatial selectivity was assessed using a modified version of the spatial information index^135^. This metric calculates the information content of firing in bits per second by comparing the firing rate in each spatial bin to the mean firing rate across the track, weighted by occupancy probability, i.e., time spent in each spatial bin. The time spent in each bin was used to normalize firing rates, and the spatial information was computed (Eq. 13) as:

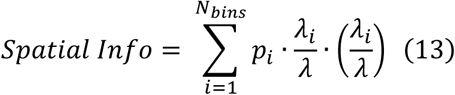

Where *p*_*i*_ is the occupancy probability of bin *i*, *λ*_*i*_ is the firing rate in that bin and *λ* is the mean firing rate across all bins, *N*_*bins*_.

### Multivariate Random Forest

To assess whether the structural features of synaptic clusters could predict the functional output of the simulated neurons, we employed a supervised machine learning approach using a Random Forest regression model. Specifically, we trained a multi-output regression model to predict place field properties (i.e., Spatial Information, In-field and Out-of-field Firing Frequency, and Field Size) from domain-specific morphological and topological synaptic features (i.e., Interspine interval std, Number of clusters, Size of clusters, Path distance of clusters, Volume of clusters, Percentage of infield branches, Average synaptic volume). These features were normalized to [0,1] range using a Min-Max Scaler to ensure comparability across dimensions. The target output matrix comprised the aforementioned spatial metrics, which were likewise scaled prior to training. For the analysis we used simulations conducted under all distribution protocols, excluding those that exhibited overflow or contained missing values.

We used a Multi-Output Regressor wrapping a Random Forest Regressor (100 estimators per model). Models were trained and evaluated over 10 independent runs using an 80/20 train-test split. Control protocols were validated against two baselines: one involving the use of random values as input features (Random training inputs, not shown) and another involving selective removal of key input features (Removal input feature, not shown) to confirm the robustness and informativeness of the cluster descriptors. Model performance was quantified using the MSE. To allow for cross-condition comparisons, the MSE for each functional output was normalized against the MSE of the "Randomized Inputs" baseline. These normalized errors were then averaged across all outputs to produce a single performance metric per run. Feature importance values were computed as the average importance across all estimators within the ensemble.

### Simulation environment and statistical analysis

All simulations were performed on our High-Performance Computing Cluster (Rocks 7.0) with 624 cores and 3.328 TB of shared RAM under CentOS 7.4 OS^136^, through the NEURON simulator^137^ using Python via an Anaconda virtual environment. Neuronal models have been simulated at a sampling rate of 40 kHz (dt = 0.025 ms). For all standard statistical tests (detailed in all relevant figure legends), the α was chosen to be 0.05 for statistical significance. To correct for multiple comparisons, the α was divided by the number of tests according to the Benjamini-Hochberg (FDR) method. Throughout the figures, p values are denoted by * (p < 0.05), ** (p < 0.01), and *** (p < 0.001).

## Supplementary figures

**Figure S1:**
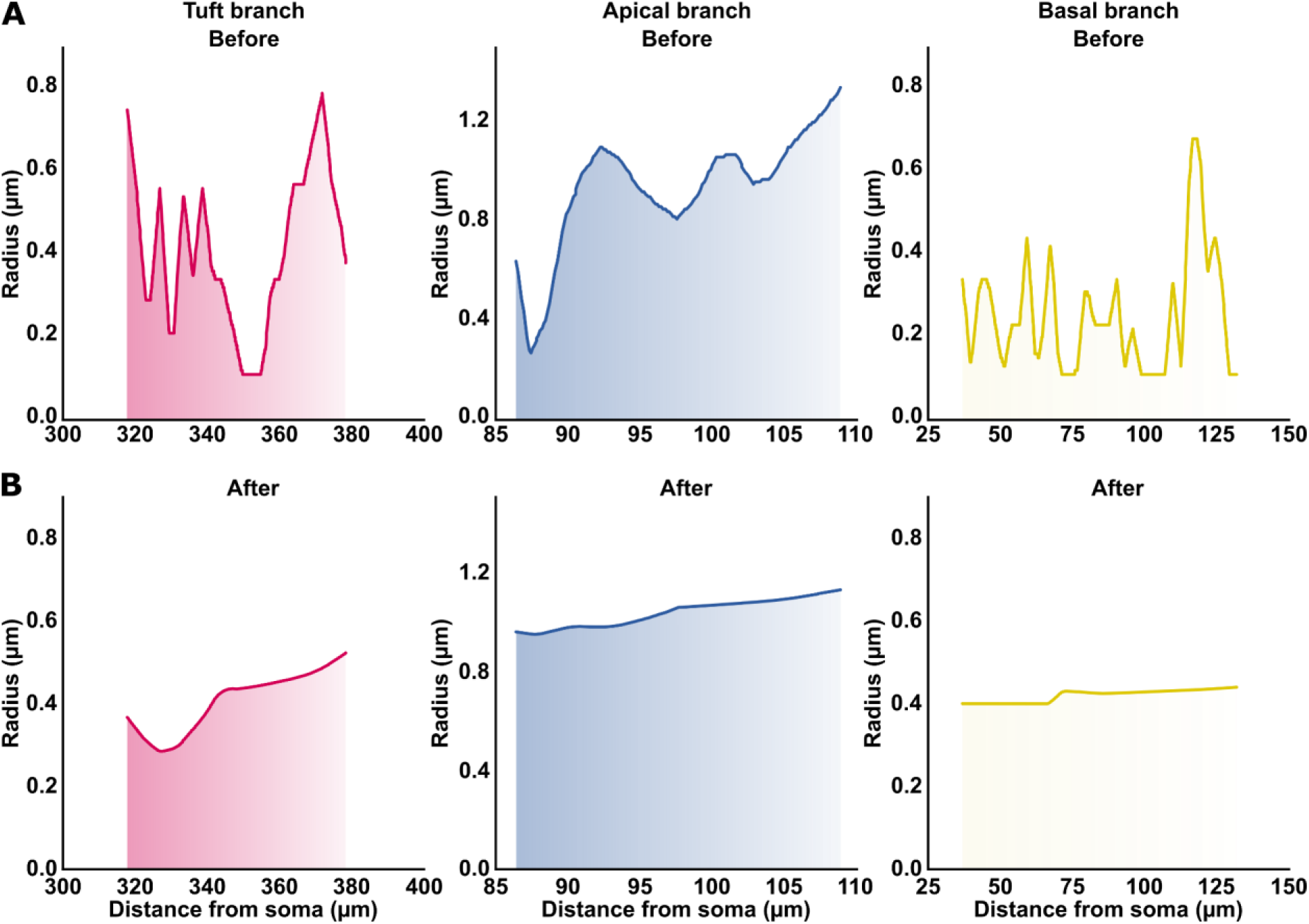
Dendritic diameter smoothing. Radius profiles of three example dendritic branches before (A) and after (B) application of the smoothing algorithm. Profiles are shown for Tuft (left), Apical (center), and Basal (right) branches, with radius plotted as a function of the distance from the soma.

**Figure S2:**
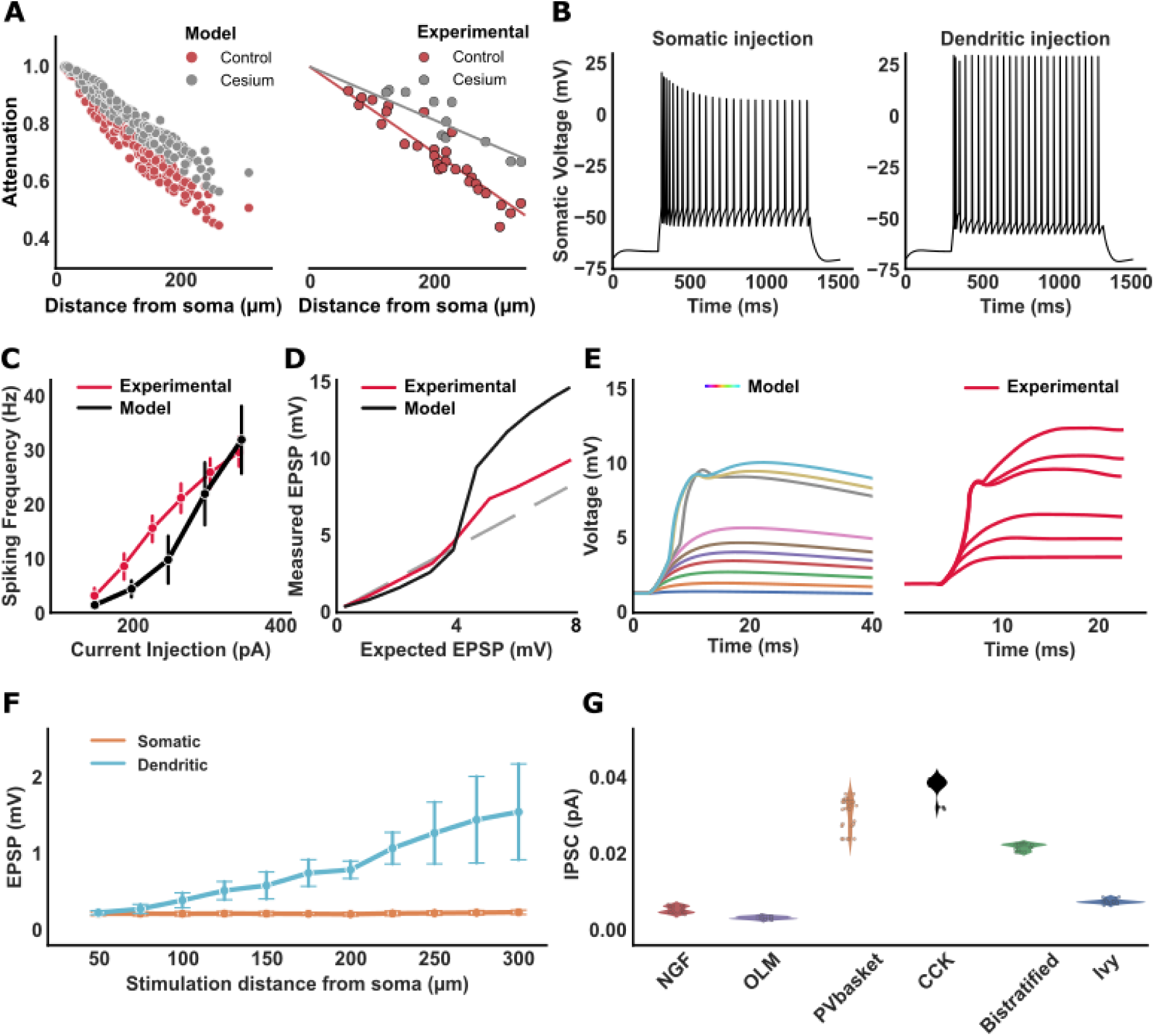
Electrophysiological Properties of CA1 PN Models Match Experimental Data. **A.** Dendritic attenuation of somatically injected negative current in control conditions (red) or simulating extracellular cesium (grey, 80% reduction of the h-current). Experimental data redrawn from Golding et al., 2005 are similar to model outputs. **B.** Examples of somatic voltage traces induced by a 300 pA current injection in the soma (left) or in a dendrite (right) located at ∼250 µm from the soma. **C.** The input-output (Frequency-current (F-I)) curves of the models resemble experimental ones from Masurkar et al., 2020^8^. **D.** Peak amplitude of EPSPs measured at the soma as a function of the expected arithmetic sum of individual EPSPs. Experimental example curve redrawn from Losonczy and Magee, 2006^24^. **E.** Somatic depolarization upon simultaneous excitation of a progressively increasing number of synapses. Left: stimulation of 1 to 30 simulated synapses on an oblique dendrite. Right: spines stimulated through glutamate uncaging, redrawn from Losonczy and Magee, 2006^24^. **F.** Dendritic and somatic excitatory postsynaptic potentials (EPSPs). Synaptic AMPA conductance strength increases with path distance from the soma, ensuring invariant somatic depolarization. **G.** Inhibitory postsynaptic currents (IPSCs) originating from various interneurons, including NGF (neurogliaform cells), OLM (oriens-lacunosum moleculare cells), PVbasket (parvalbumin-expressing basket cells), CCKbasket (cholecystokinin-expressing cells), Bistratified and Ivy cells. Validated to match experimentally observed data. Errorbars in **C**, and **F** represent 95% confidence interval (CI) of the mean.

**Figure S3:**
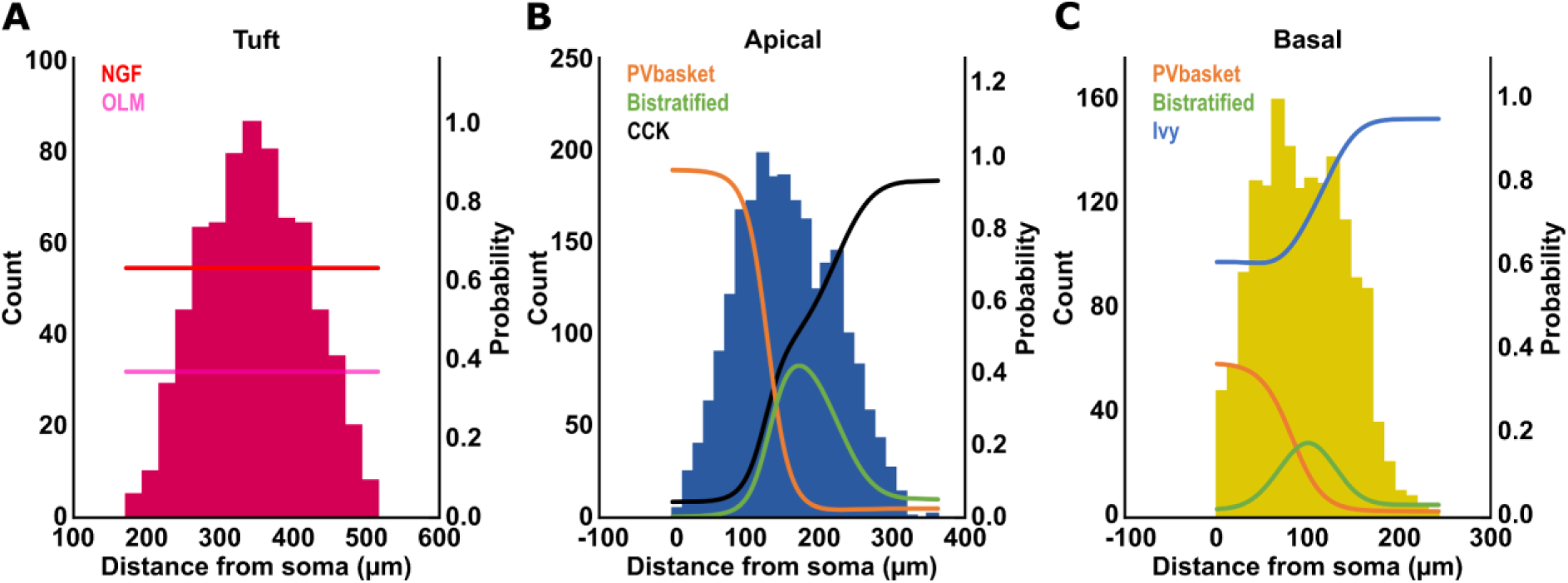
Probability of inhibitory input sources targeting synapses in CA1 pyramidal cells. Distribution of inhibitory presynaptic input probabilities for synapses located on the Tuft (A), Apical (B), and Basal (C) dendrites. The probability varies with dendro-somatic path distance, capturing how different interneuron types preferentially target specific dendritic regions. Histograms show the number of synapses as a function of distance from the soma, while the overlaid curves represent the probability that a synapse is targeted by each interneuron type.

**Figure S4:**
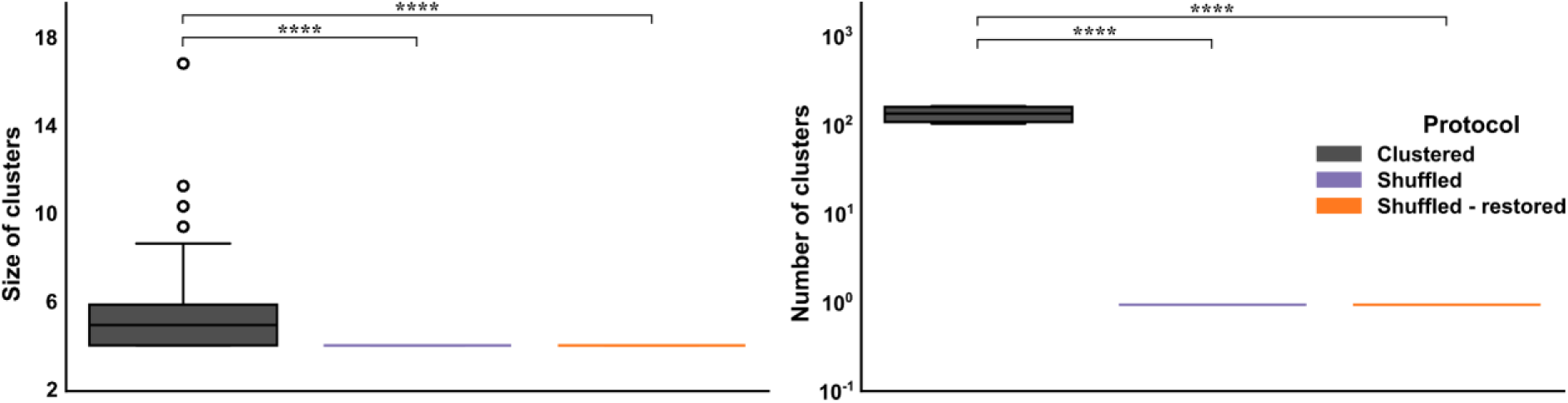
Characterization of synaptic clusters under alternative input allocation protocols. The cluster size of cotuned synapses (A) and their number (B) were significantly reduced following synaptic shuffling. Statistically significant differences were observed between the control configuration and the shuffled conditions, with variation maintained across dendritic domains. Pairwise Mann-Whitney U tests with Benjamini-Hochberg (FDR) correction were used. Significance: ∗p < 0.05, ∗∗p < 0.005, ∗∗∗p < 0.001, and ∗∗∗∗p < 0.0001. Error-bands in C, F, and G represent 95% confidence interval (CI) of the mean.

**Table S1:**
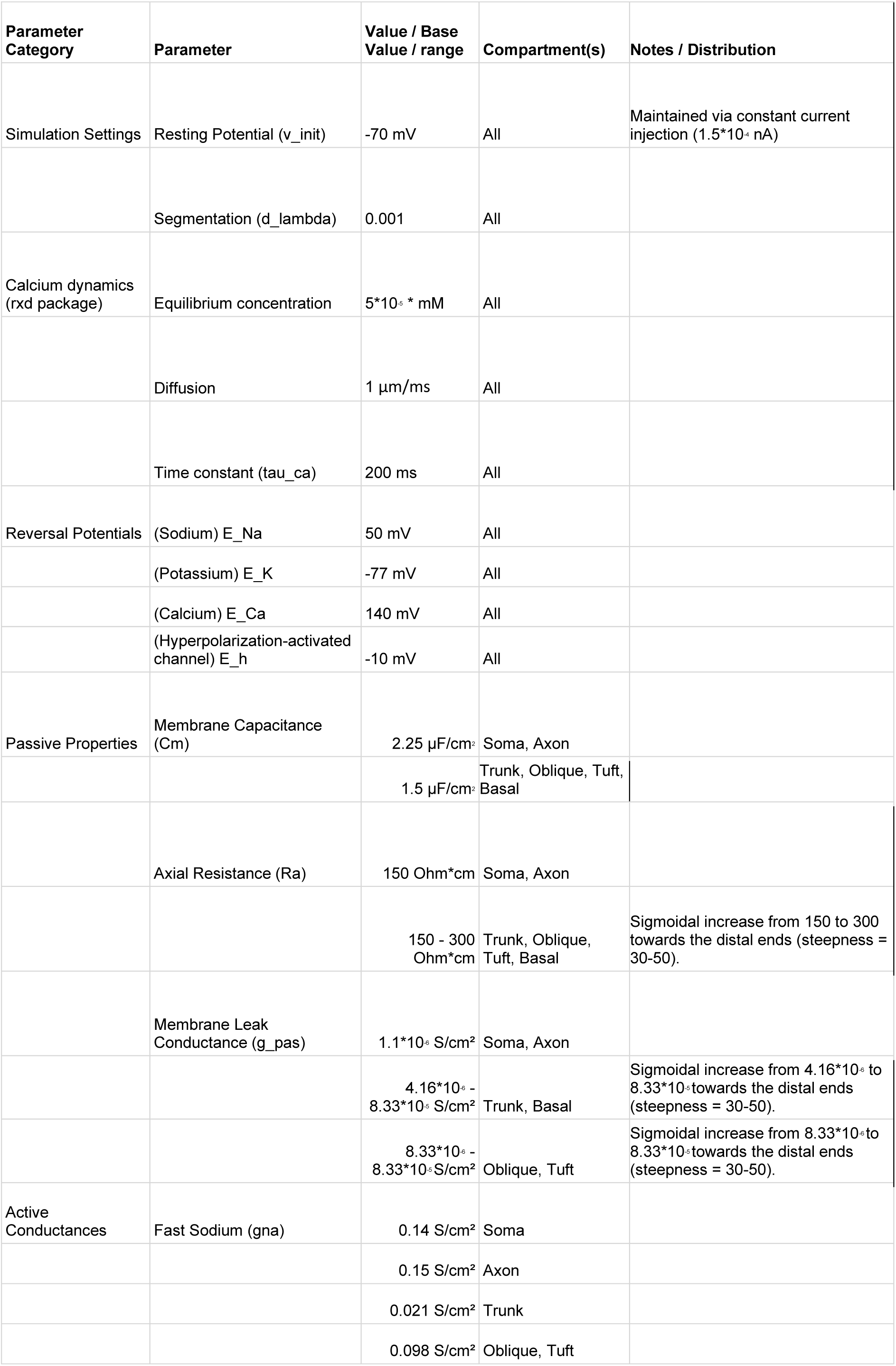
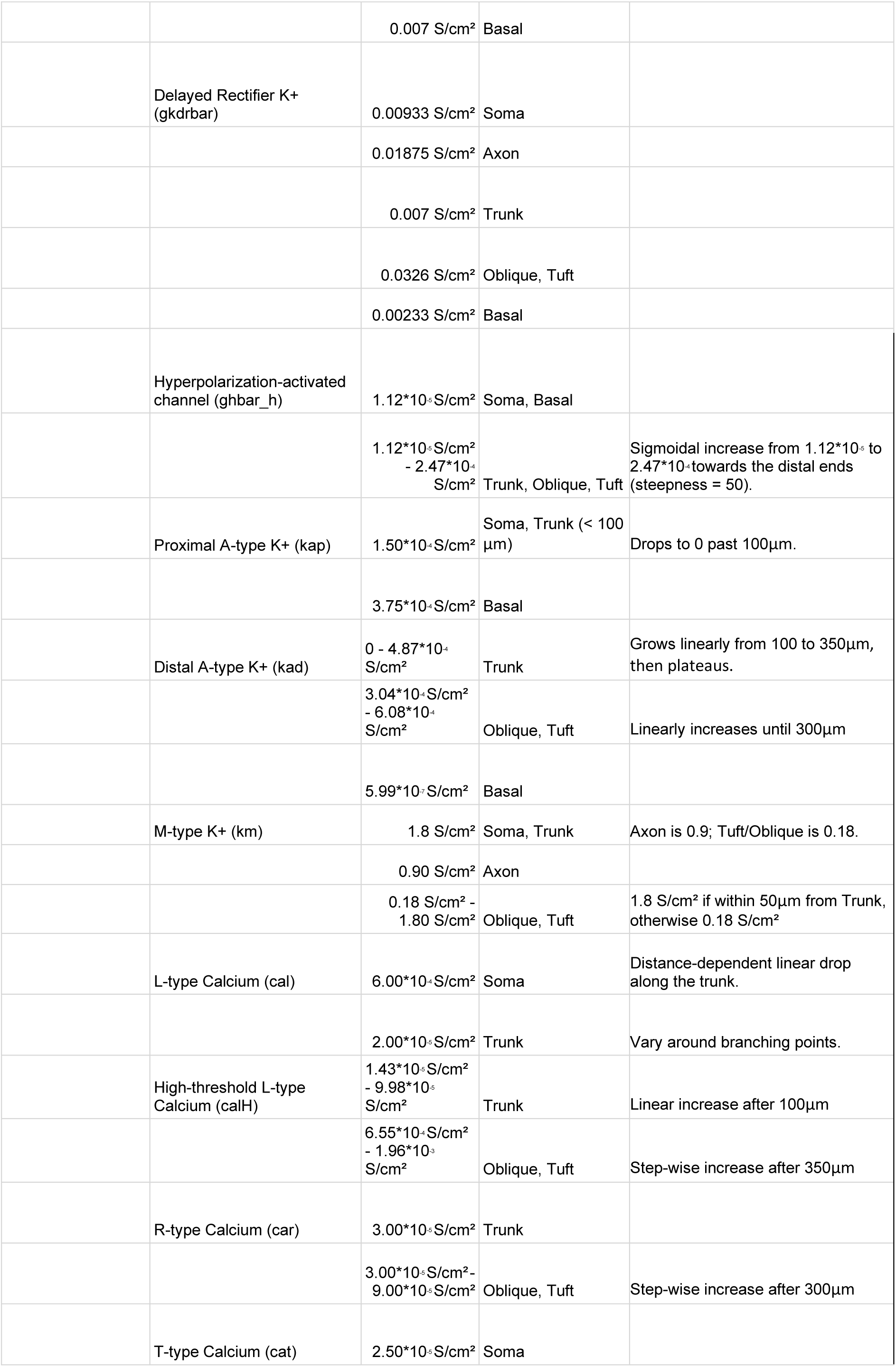
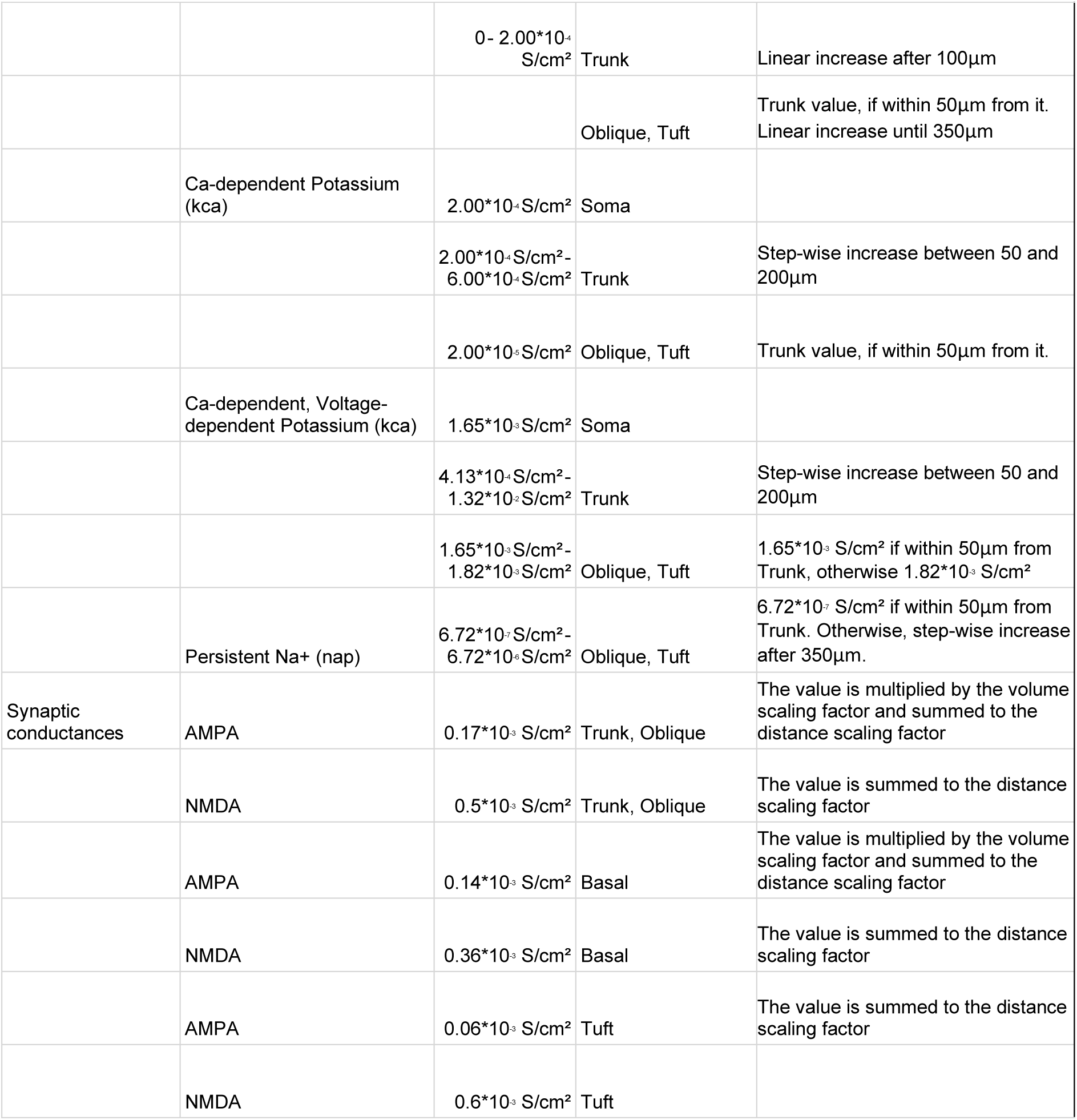
Channel hyperparameters table. The table contains the values related to the initialization settings, calcium dynamics, passive and active mechanisms, and synaptic conductances of the simulated cells. Given the complexity of the models, this table doesn’t contain the values of the channel’s kinetic and cannot give an exhaustive description of their non-linear distribution. For further details, please refer to the code.

## Supplementary Materials and Methods

### Somato-dendritic Parameter Distributions

To accurately capture the location-dependent integration properties of CA1 pyramidal neurons, several passive properties and active conductances were distributed non-uniformly across the somatodendritic axis. The model implements these gradients as a function of the path distance from the soma. To maintain consistent electrophysiological profiles regardless of specific morphological variations, distance-dependent rules are normalized to the maximum path length of the apical tree in the reference model (Poirazi et al 2003).

The scaling factor *S*_*domain*_ is defined as:

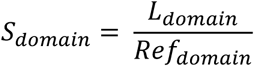

where *L*_*domain*_ is the maximum path distance (in μm) from the soma to the distal tip of the apical dendrites for a given cellular morphology, and *Ref*_*domain*_ represents the reference length. Specifically, *Ref*_*apic*_ = 447.01μm and *Ref*_*tuft*_ = 619.53μm.

### Sigmoidal Gradients for Passive Properties and h-current

In our model, several membrane properties or conductances (e.g. *R*_*m*_, *R*_*a*_, *I*_*h*_ etc.) increase or decrease along the proximo-distal axis of CA1 pyramidal cells. To model the smooth transition of these properties along the dendritic domains, we utilized a sigmoidal function:

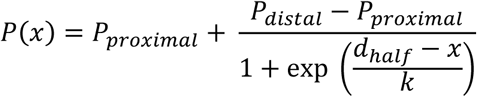

where *P*(*x*) is the parameter value at path distance *x* from the soma, *P*_*proximal*_ and *P*_*distal*_ are the baseline and asymptotic end values, *d*_*half*_ is the distance of the half-maximal inflection point, and *k* dictates the steepness of the gradient. The specific constants applied are detailed in Supplementary Table S1.

### Step Functions in Distal Compartments

To account for functional differentiation in the terminal tuft and oblique dendrites, discrete thresholds dictate the conductances of R-type calcium (*g*_*car*_) and persistent sodium (*g*_*nap*_) channels.

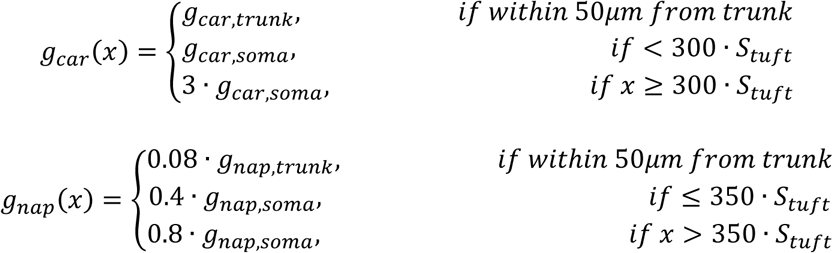

Other channels with step function conductance determination are the voltage-gated sodium (*g*_*nas*_), the delayed rectifier potassium (*g*_*kvs*_), the A-type potassium (*g*_*kap*_), the slow calcium-dependent potassium current (*g*_*kca*_), and the slowly activating muscarinic potassium (*g*_*km*_) channels.

### Piecewise Linear Gradients for Active Conductances

Several voltage-gated channels on the apical trunk exhibit linear distance-dependent scaling up to specific saturation points, facilitating distal dendritic excitability and precise spike backpropagation.

For example, the distal A-type potassium conductance (*g*_*kad*_) increases linearly starting at ∼100μm from the soma:

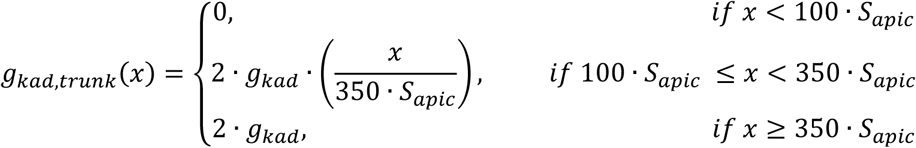

Similarly, low-voltage-activated T-type (*g*_*cat*_) and L-type high-voltage-activated (*g*_*calH*_) calcium conductances scale linearly to support the generation of distal dendritic calcium spikes:

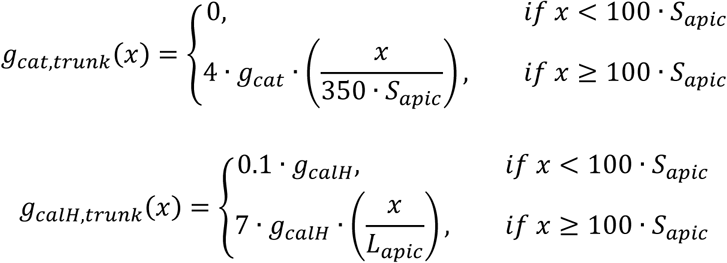

More conductances present a linear gradient progression, like the L-type calcium channel with low threshold for activation (*g*_*cal*_), and the T-type calcium channel with a high threshold for activation (*g*_*cat*_).

The equations reported here represent only a few examples. For more detail, refer to the code.

